# HIV-Attributed Causes of Death in the Medical Ward at the Chris Hani Baragwanath Hospital, South Africa

**DOI:** 10.1101/388462

**Authors:** Andrew Black, Freddy Sitas, Trust Chibrawara, Zoe Gill, Mmamapudi Kubanje, Brian Williams

**Affiliations:** Department of Internal Medicine, University of the Witwatersrand, Johannesburg, South Africa; Menzies Centre for Health Policy, Sydney School of Public Health, University of Sydney, Camperdown and School of Public Health and Community Medicine, University of New South Wales, Kensington, Australia; South African Centre for Epidemiological Modelling and Analysis, Stellenbosch University, Stellenbosch, South Africa

## Abstract

**Background:** There are sparse data in Africa on the association between HIV infection and deaths from underlying medical conditions. Using records from the Chris Hani Baragwanath Hospital (CHBH) in Soweto, South Africa, we determined mortality from medical conditions associated with HIV.

**Methods:** From January 2006 to December 2009 AB collected data on 15,725 deaths including age, sex, day of admittance and death, HIV status, ART initiation and CD4+ cell counts and reviewed the underlying cause of death using medical notes. Conditions known to be associated with HIV were cases; conditions not associated with HIV were controls. We calculate the HIV odds-ratios for cases relative to controls and HIV-attributable deaths as the fraction of those with each condition, the disease-attributable fraction (DA), and as the fraction of all deaths, the population-attributable fraction (PAF).

**Interpretation:** The high prevalence of HIV among those that died in the medical wards at the CHBH, especially in those below the age of 50 years, demonstrates the impact of the HIV-epidemic on adult mortality and hospital services and the extent to which early antiretroviral treatment would have reduced the burden of both. Of the deaths included in the analysis the prevalence of HIV was 61% and the prevalence of AIDS related conditions was 69%. The HIV-attributable fraction was 36% in the whole sample and 60% in those that were HIV-positive. Cryptococcosis, Kaposi’s sarcoma and *Pneumocystis jeroveci* are highly predictive of HIV while TB, gastroenteritis and anaemia are very strongly associated with HIV. The greatest number of deaths attributable to HIV was among those dying of TB or of other respiratory conditions.

**Funding:** No funding was received for this study.

## Research in Context

### Evidence before this study

A study using data collected between 2002 and 2006, immediately before these data were collected, suggested that 1.8% of deaths in a public hospital in the Eastern Cape, South Africa were due to HIV.^1^ However, the HIV-status of those that died was not recorded showing the importance of being able to estimate the proportion of deaths due to particular conditions that can be attributed to HIV infection. A more recent study^2^ suggested that in 2006 283k deaths or 42% of all deaths in South Africa were attributable to HIV, and the authors compared this to other estimates for 2006 of 225k using the ASSA Model,^3^ 250k using the Thembisa model,^2^ 270k in the Global Burden of Disease study,^4,5^ 350k using the UNAIDS Spectrum/EPP model and 354k based on changes in the age-distribution of deaths over time.^6^ The considerable variation in these estimates was attributed to misclassification of AIDS deaths associated with different underlying conditions.^2^

The difficulty in estimating AIDS deaths is partly due to the difficulty of deciding what proportion of deaths from a particular cause, such as TB, should be attributed to HIV.^7^ The published models^2–8^ all agree that the number of AIDS deaths peaked in 2006 but the model estimates of mortality vary widely because the models differ in their assumptions about survival after infection without ART. Without a direct estimate of AIDS-related deaths or of survival after infection without ART, models fitted to trend data in HIV prevalence and ART coverage cannot be used to accurately determine the overall mortality attributable to HIV.

### Added value of this study

By reviewing and assigning underlying causes of death to their appropriate categories we derive more accurate estimates of their burden in relation to HIV which are important for politicians, clinicians, the health system, and for modelling the epidemic. The present study was completed just as the roll-out of ART started in the public health services and provides a snapshot of the HIV-burden among those who were not on ART. Now that about two-thirds of people infected with HIV are on ART^9^ this should greatly reduce AIDS-related mortality and the data presented here provide a baseline against which future studies can be compared.

### Implications of all the available evidence

While several studies have used indirect methods to assess the disease attributable-fraction (DAF) and the population-attributable fraction (PAF) of deaths attributable to HIV, this is the first study in South Africa to determine these fractions directly using data on the cause of death combined with data on the HIV status of those that died at a time when ART treatment was largely unavailable. This will help to put estimates of the number of deaths that can be attributed to HIV on a firm footing. Now that ART has been rolled out more widely a similar study, including data on the ART status of those that died, would show directly the impact of ART on mortality in South Africa.

## Introduction

The early response of the South African Government to the epidemic of HIV was one of confused denial^10^ and although the epidemic started later in South Africa than in many neighbouring countries^9^,^11^ the provision of ART in the public health system only began in earnest in 2006 by which time AIDS deaths had reached their peak.^9^ The stigma associated with HIV and the unwillingness to acknowledge the magnitude of the epidemic meant that HIV and AIDS were often omitted from death certificates with other causes of death, such as TB, listed as the preferred alternative.

Between 2006 and 2009, when these data were collected, the Chris Hani Baragwanath Hospital (CHBH) was one of the main referral hospitals in South Africa serving a population of 1.3 million or 2.4% of the population of South Africa. At that time about 300 thousand people were dying of AIDS each year^2^ so that about seven thousand would have died each year in Soweto. In this study there 3.9 thousand deaths each year, on average, suggesting that more than half of all those that died of AIDS in Soweto died in the medical wards of the CHBH. In an earlier study among children in the CHBH^12^ the proportion of deaths accounted for by HIV infection increased from 7% in 1992 to 46% in 1996 with an overall odds ratio of 2.85.

Here we use a detailed and extensive set of data from the CHBH where the cause of death as well as the HIV-status of those that died was established in order to determine the proportion of deaths from a range of causes that are attributable to HIV before ART became widely available in the public health sector.

## Methods

### Study design

The study was conducted in the medical ward of the CHBH. The trauma, emergency, obstetrics, gynaecology, surgery, and paediatric wards were not included. For all 15,725 adults that died between January 2006 and December 2009 the hospital number, age, sex, cause of death, date of admission, date of death, HIV status, CD4+ cell count, and ART status were recorded on the Baragwanath Mortality Record (BMR). Owing to the stigma associated with HIV, clinicians were reluctant to include HIV status or AIDS defining causes of death on the official death notification forms. The BMR was developed to allow for a more accurate and detailed record of the causes of death. Attending medical consultants ascertained the underlying causes of death by reviewing the patient medical notes and completed the BMR at the time of signing the deceased’s official death certificate. Data from the BMR were entered into an Excel spread sheet and the cause of death was classified according to ICD 10 by a trained data capturer. In cases where the code allocation was unclear to the data capturer AB reviewed the available clinical data and assigned the ICD 10 code.

### Data cleaning

The data are missing 9 hospital numbers, 109 dates of admission, 1 date of death, no ages or sex and 529 ICD codes that make up 3.4% of the sample (Supporting Information, Appendix 1 and 2). In the whole sample the HIV status of 11% was given as ‘suspected’ and of 15% as ‘unknown’.

### Grouping causes of death

There were 323 individual ICD codes in the data and the number of deaths, HIV-status, mean age, number of men and women, and the groups to which they are assigned are given in the Supporting Information (Appendix 2). For 529 deaths (3.4% of the total sample) an ICD code was not assigned and there was uncertainty as to whether or not 527 poorly defined deaths (3.4% of the sample) were from conditions associated with HIV. The remaining 14,669 deaths were assigned to one of 15 conditions associated with HIV/AIDS or to a control group.

We carried out a case control study to measure the HIV-attributable fraction in cases as compared to controls. Cases were the following AIDS related conditions with their ICD-10 codes: infectious gastroenteritis (A09), tuberculosis of the lung (A15 and A16), extra-pulmonary tuberculosis (A17 to A19 and A31), certain infectious and parasitic diseases (A41), HIV (B21 to B24), Cryptococcosis (B45), pneumocystis (B59), Kaposi’s sarcoma (C46), Hodgkin’s and non-Hodgkin’s lymphoma (C82 and C85), diseases of the blood and blood forming organisms (most of D51 to D89), meningitis (G00, G03, G04, G06 and G08), pneumonia, chronic obstructive pulmonary disease, and lower respiratory tract infections (included in J01-J99), diseases of the digestive system (K72-K74), diseases of skin and bone (included in L03-L95 and M01-M99) and genitourinary conditions (N00-N39). Controls were medical conditions that are not, or are not known to be, associated with HIV and included: malignant neoplasms excluding Kaposi’s sarcoma, Hodgkin’s and non-Hodgkin’s lymphomas, disorders involving immune mechanisms or the nervous system, heart disease and stroke, and external causes of death such as injury or poisoning (details in the Supporting Information, Appendix 2).

### Statistical Analysis

We first consider the overall effect of age and gender on the number of people that died by fitting the number of deaths to skew-normal distributions (Supporting Information, Appendix 3). Because some deaths were given a ‘suspected’ or ‘unknown’ HIV status, we use the fitted curves to estimate the proportion of those whose HIV-status was ‘suspected’ or ‘unknown’ that were in fact HIV-positive or HIV-negative and use this to justify assigning HIV-suspected cases to the HIV-positive category and HIV-unknown cases to the HIV-negative category (Supporting Information, Appendix 3). We then carried out a case-control analysis, adjusting for age, to determine the odds-ratio (OR) for HIV in each of the AIDS related conditions as compared to the control conditions and to calculate disease and population fractions attributable to HIV (Supporting Information, Appendix 4).

## Results

For HIV-negative men and women the modal ages at death were 64 and 74 years, respectively; for HIV-positive men and women they were 37 and 32 years, respectively (Figure 1). CD4^+^ cell counts were available for 41% of those that were HIV-positive and among these the median CD4+ cell count was 45/μL (90% between 3/μL and 310/μL) showing that most of the HIV-positive patients were in late stages of HIV. Despite the availability of ART in the public sector ART coverage was low in those infected with HIV increasing from 4.9% in 2006 to 8.6% in 2009.

**Figure 1.**
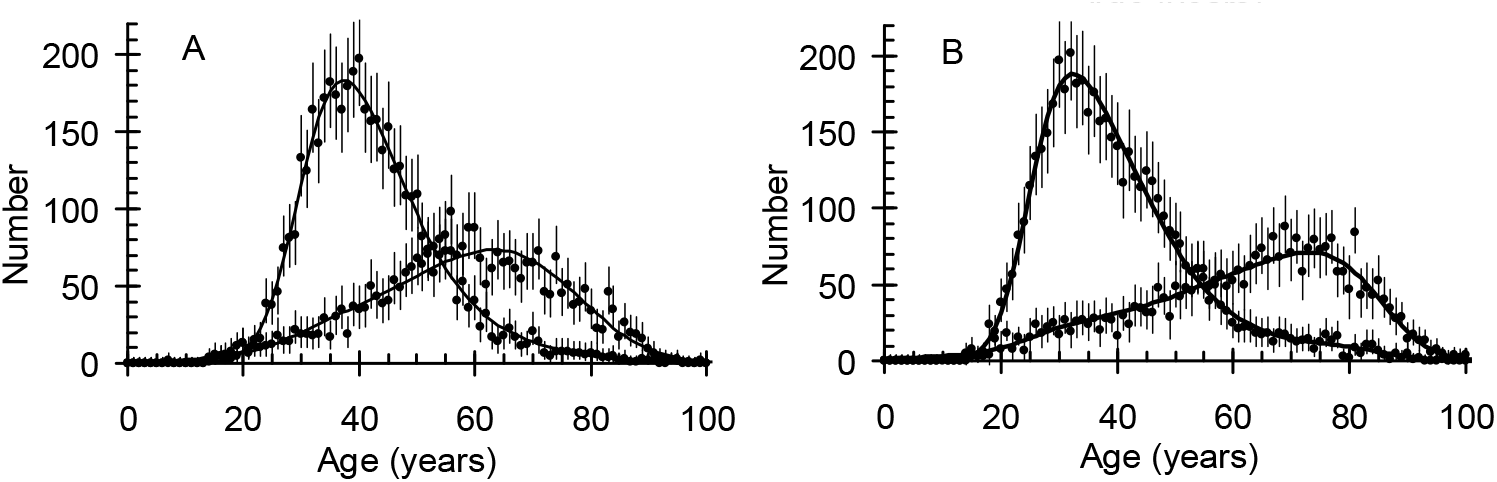
The number of A: Men and B: Women who were HIV-positive (peaks at lower ages) and were HIV-negative (peaks at higher ages).

The distribution of time from admission to death was not significantly different for men and women or for those that were HIV-positive or HIV-negative. Amongst this deceased population, 27% died by the day after admission and the mortality dropped to 11% to 13% per day after that (Supporting Information, Appendix 5).

## Prevalence of HIV in controls

The prevalence of HIV in those with the control conditions is given in Figure 2 and the Supporting Information (Appendix 6). The peak prevalence is high at about 67% in 25-35 year olds and even in those aged 80 to 90 years the prevalence of HIV is 12% ± 3%. Comparing the prevalence of HIV in the control group to the prevalence of HIV in the antenatal clinic surveys in the Johannesburg Municipality from 2006 to 2009,^13^ where the peak prevalence was 40%, suggests an odds ratio of 3.2 but the overall distributions of deaths by age are closely matched. We discuss the reasons for this difference and the implications for this analysis in the Supporting Information (Appendix 6).

**Figure 2.**
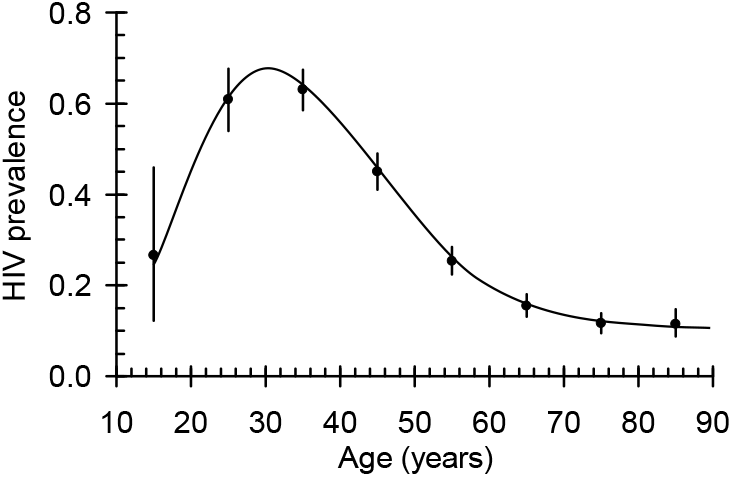
The prevalence of HIV in the control group as a function of age.

### Note: 1

#### Odds and odds ratios

The odds for being HIV positive given each condition and the odds ratios compared to the control group were calculated in ten-year age bands (Supporting Information, Appendix 7). The adjusted odds ratios are the weighted averages over all ages. For four of the conditions, cryptococcosis, Kaposi’s sarcoma, *Pneumocystis jeroveci,* and diseases of the blood and blood forming organs, there were too few negative cases to adjust reliably for age. We therefore established a relationship between the adjusted and the crude odds ratios (Supporting Information, Appendix 8) and used this to estimate the age-adjusted odds ratios for these four conditions.

The final estimates for the ORs, DAFs and PAFs are shown in Table 1 and Figure 3 and Figure 4. For cryptococcosis, Kaposi’s sarcoma, pneumocystis and tuberculosis the DAFs are range from 80% to almost 100%. For gastroenteritis, anaemia, meningitis and lymphoma the DAF range from 50% to 75%. For other respiratory conditions, sepsis, genitourinary and disorders of the skin and bone the DAF ranges from 30% to 50%. For digestive conditions the DAF is 16%.

**Figure 3.**
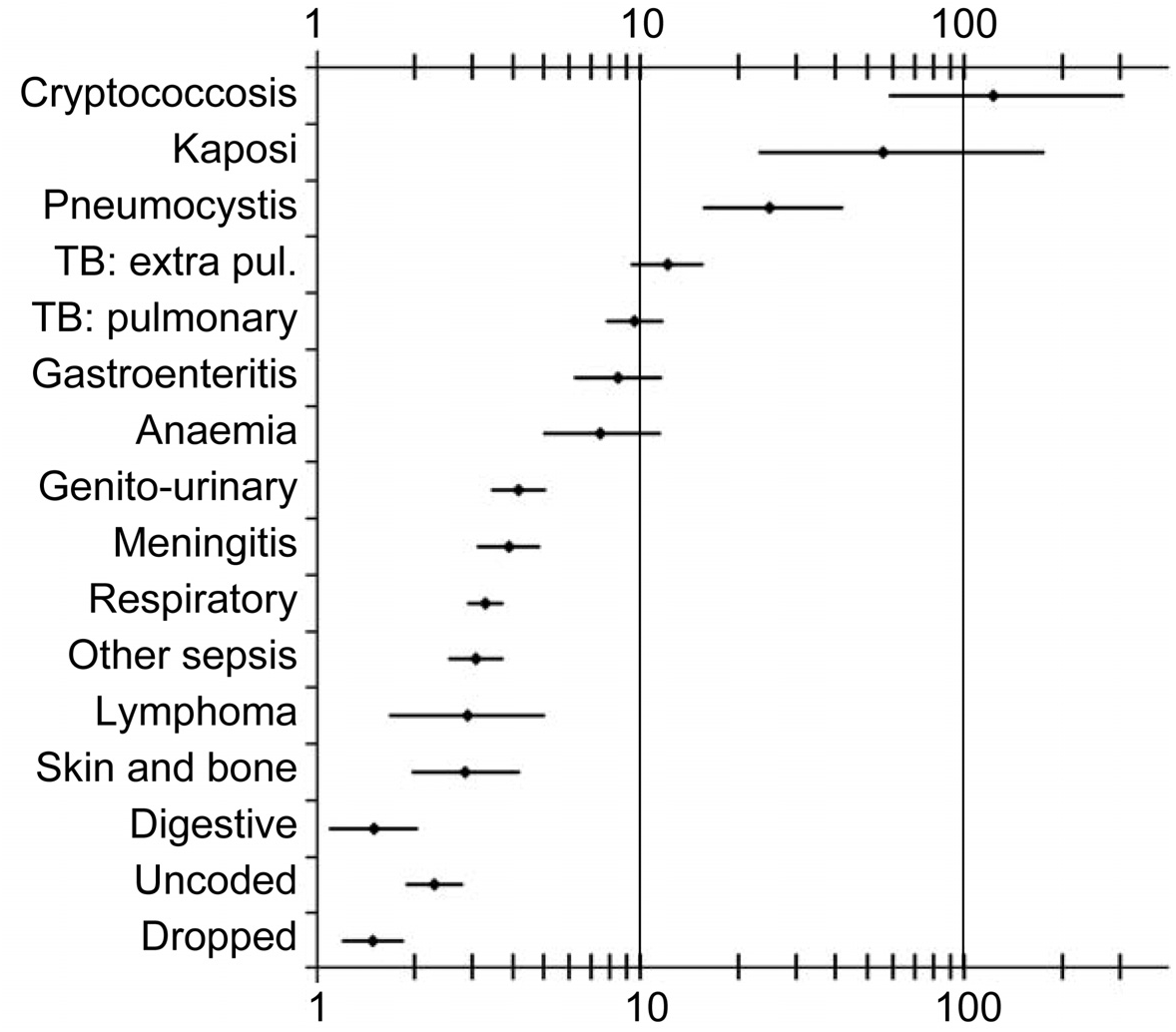
HIV-odds ratios for causes of death compared to controls.

**Figure 4.**
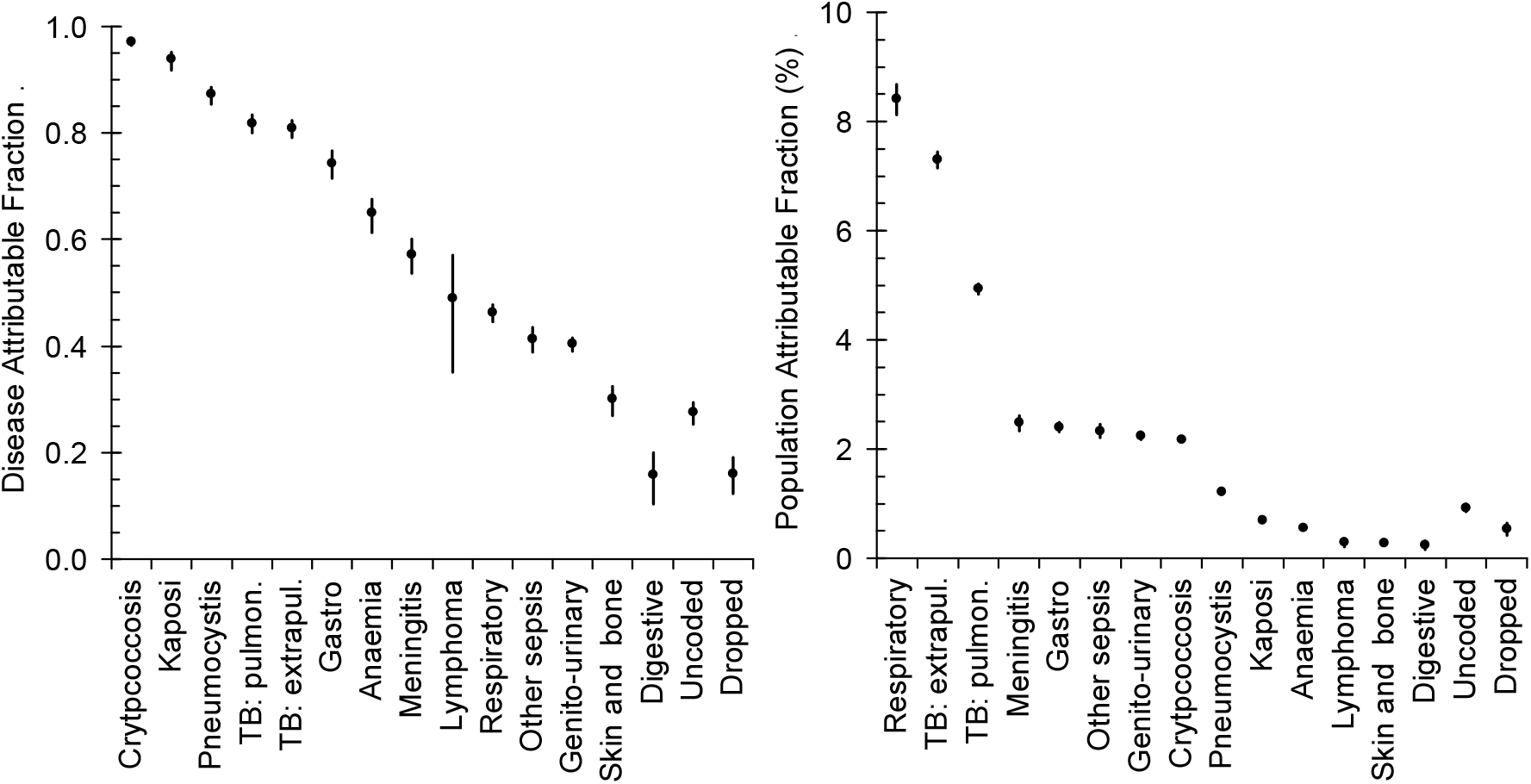
HIV disease-attributable fraction (DAF) and population-attributable fraction (PAF) for different causes of death.

**Table 1.**
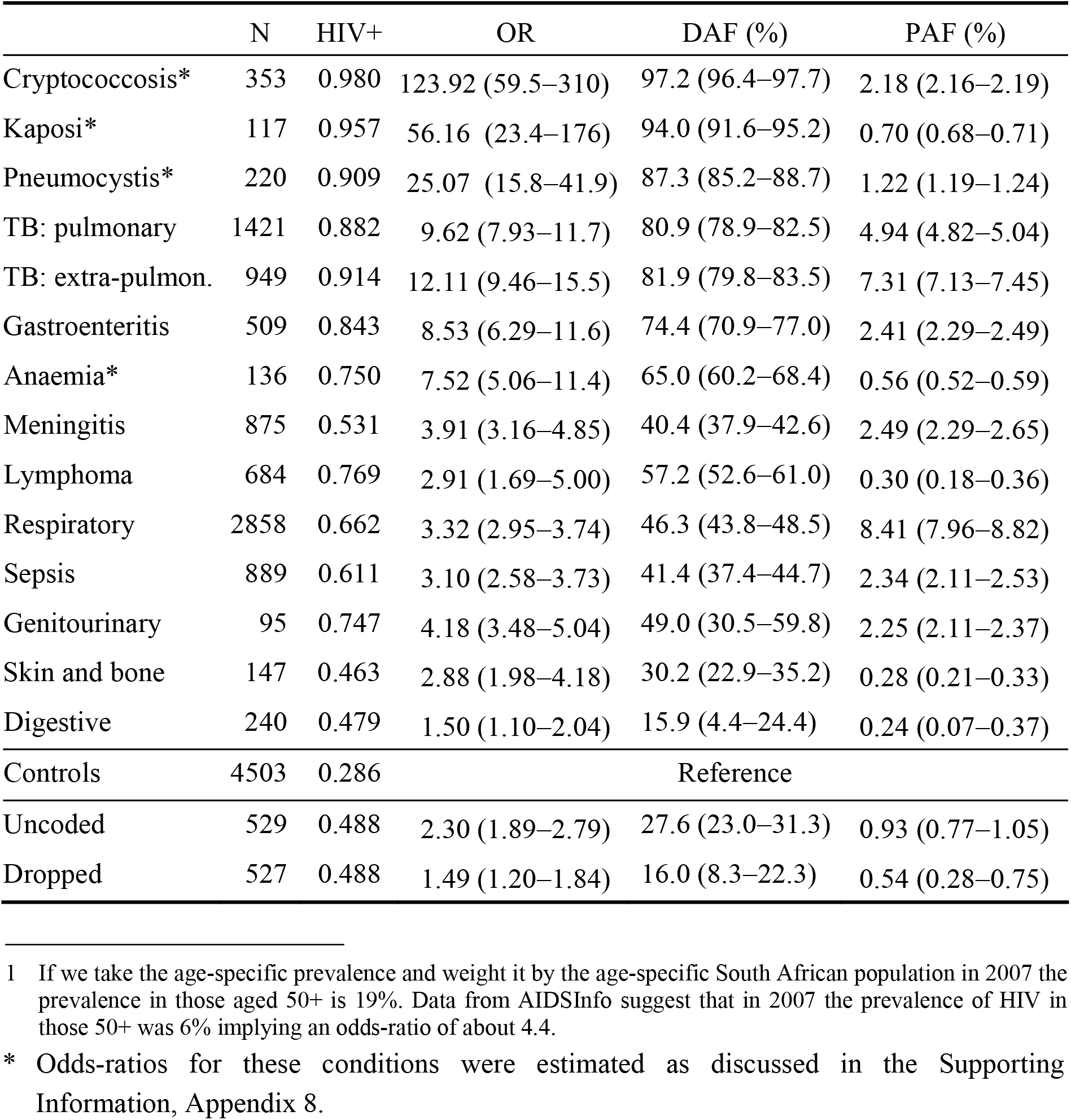
The total number, prevalence of HIV, odds ratios for causes of death versus control conditions not associated with HIV, disease (DAF) and population (PAF) attributable fractions. Point estimates with 95% confidence limits. (Details in Supporting Information, Appendix 7.)

The PAF is given by the DAF multiplied by the prevalence of that condition (Supporting Information, Appendix 4). HIV-attributable deaths from TB and other respiratory conditions together account for 20.7% ± 0.9% of all deaths. HIV-attributable deaths from cryptococcus, gastroenteritis, meningitis, sepsis and genitourinary conditions attributable to HIV each account for between 2% and 3% of all deaths and together they account for 11.7% ± 0.6% of all deaths. HIV-attributable deaths from Kaposi’s sarcoma, pneumocystis, anaemia, lymphoma, diseases of the skin and bone and of the digestive system each account for less than 1.5% of all deaths and together account for 3.3% ± 0.4% of all deaths.

Of the deaths included in the analysis, excluding those that were not coded or were dropped, the prevalence of HIV was 61% and the prevalence of AIDS related conditions was 69% (Table 2). The HIV-attributable fraction in the whole sample was 36% and in those that were HIV-positive the HIV-attributable fraction was 60%. In 2006 the estimated number of deaths in South Africa was 691k in a study by Bradshaw *et al.^2^* and 738k in a study by UNAIDS.^8^ If we take the average, 706k deaths, this study would suggest that the HIV-attributable deaths were about 254k, close to the estimate using the Thembisa model.^2^

**Table 2.** The total number of deaths included in the analysis, the number that were HIV-positive, the number that had an AIDS-related condition (ARC), the number attributable to HIV, the prevalence of HIV and AIDS related conditions, the proportion of deaths attributable to HIV among all deaths and among those that were HIV positive.

|  |  |  |  |
| --- | --- | --- | --- |
| All deaths | 14,669 | HIV prevalence | $61\% \pm 1\%$ |
| HIV+ | 8,935 | ARC prevalence | $69\% \pm 1\%$ |
| ARC | 10,166 | Attributable/Total | $36\% \pm 1\%$ |
| HIV attributable | 5,321 | Attributable/HIV-positive | $60\% \pm 2\%$ |

## Discussion

We estimate the excess mortality from conditions known, or suspected from the literature, to be HIV/AIDS related. Our imputation of HIV positive and negative patients goes some way to avoiding classifying deaths by their exposure status bearing in mind that in some cases HIV/AIDS is given as the cause of death. The data in Table 1 show that almost all deaths from Kaposi sarcoma, cryptococcosis or *Pneumocystisjeroveci* infections could be attributed to HIV. Kaposi sarcoma was rare before the HIV epidemic struck^14^ and odds ratios of 56 (23-176) and a DAF of 94% (92%-95%) concur with previous work in Soweto, Johannesburg in which the corresponding OR versus controls was 47 (32-70)^15^ while 89% of patients with Kaposi sarcoma were HIV positive. Likewise cryptococcosis or *Pneumocystis jeroveci* infections were extremely rare pre-HIV, only occurring in persons who were immune compromised.^16^ ORs for lymphomas in this analysis of 2.9 (1.7-5.0) resemble those found in a previous case control study where the OR was 5.9 (4.3-8.1) for non-Hodgkin and 1.6 (1.0-2.7) for Hodgkin lymphoma.^15^

The prevalence of HIV in the control group is high and comparing this to the prevalence of HIV in ante-natal clinic surveys between 2006 and 2009 in the Johannesburg Municipality suggests an OR of 3.2. However, the prevalence of HIV among pregnant women in the Johannesburg Municipality may have been lower than in Soweto and it could also be that HIV causes such a force of mortality that even for deaths that are unrelated to HIV, such as car accidents or lung cancer, being HIV positive would cause additional medical complications leading to a greater case fatality. The high prevalence of HIV in older people has been acknowledged^14^ but only touched on briefly in the literature and is particularly striking since the life expectancy without ART of those aged 85 years is only about 2 years.^17^ However, a high prevalence of HIV of 2.6% in those aged 65 years or more was found in control arm of a study of newly diagnosed cancers at CHBH and Johannesburg Hospitals between 1995 and June 2004.^18^ Various explanations have been put forward^12^ but the reason for this high prevalence in older people remains unclear. As shown in the Supporting Information (Appendix 6) increasing the odds-ratios by a factor of 3.2 increases both the DAFs and the PAFs but the overall pattern and the general conclusions remain unchanged.

Even though the deaths examined did not cover all wards and all hospital deaths, such as deaths from cancer of the cervix, an AIDS-defining condition, and a mixture of deaths attending the intensive care unit, it is clear that the proportion of deaths in this series that are attributed to HIV are high, ranging from 20% of deaths from digestive causes to almost 100% for Kaposi sarcoma, *Pneumocystis jeroveci* and cryptococcal infection.

Tuberculosis, for which the PAF due to HIV is in excess of 90% is the most common opportunistic infection among people infected with HIV^18–20^ and TB co-morbidity is very likely in Soweto where tuberculosis is endemic. Specific diagnoses of the organisms underlying meningitis, gastroenteritis and respiratory conditions were not always available and a correct diagnosis requires a culture which is costly and time consuming.^21^ In sub-Saharan Africa, there is limited literature on the causative agents of meningitis, gastroenteritis and other infection related deaths but the aetiology of these is different in those with and without HIV.^21^

There is evidence that anti-retroviral therapy causes a decline in the incidence of several AIDS defining conditions, most clearly in the case of TB,^22^ but full immune recovery is only achieved if treatment is started immediately after infection when CD4^+^ cell counts are still high.^23^ Of all the deaths between 2006 and 2009 in the medical ward of CHBH up to 36% could have been averted (Table 2) if people living with HIV had been started on antiretroviral therapy early in the course of their infection. In countries with well-established treatment programs, the life expectancy for people with HIV approaches that seen in HIV-negative people^24^ but in 2006-2009 the policy was not to start people on ART until their CD4+ cell-count had fallen to less than 200/μL^15^,^18^ by which time their immune system was severely compromised. In this study 6.3% of those that were HIV-positive were on ART which matches the 6% coverage of ART among South African adults at that time.^9^ The mean age at death for those infected with HIV was 38 years for men and 33 years for women, suggesting that millions of life-years were lost due to the low coverage and late provision of treatment. Providing ART much earlier in the epidemic would have saved the lives of millions of young adults while minimizing the economic impact of HIV, and reducing the burden on the health system.^25^ A recent study of productivity losses due to premature mortality from cancer showed that the cost per cancer death in South Africa was US$101k. The 82 deaths from Kaposi’s sarcoma alone will have cost US$8.3M in lost productivity.^26^ If we estimate the cost of keeping one person in CHBH for one day at about US$100 and noting that the average survival after admission was 6.7 days, then the hospital costs of the 2.5k HIV-attributable deaths each year will have amounted to US$1.7M*per annum.*

It will be of great importance to repeat this study in the same or a similar hospital as this will provide a direct estimate of the impact of the roll-out of ART on mortality from AIDS related conditions in South Africa.

## Contributors

AB carried out the study and collected the data. All authors contributed to the analysis and interpretation of the data and writing the paper.

## Ethical approval

The University of the Witwatersrand Human Research Ethics Committee gave ethical approval (Number M111103).

## Supplementary Material

### Appendix 1: Data cleaning

#### Dataset

Data were captured using a datasheet developed by AB in 2006 and referred to as the Baragwanath Mortality Record (BMR). Medical consultants completed the datasheet at the time of signing the deceased’s official death certificate. In some cases, the underlying cause of death reported on the BMR datasheet differed from that recorded on the official death notification form. In such cases, the cause of death was ascertained by reviewing the patient file. Data were captured in a Microsoft Excel Workbook with the cause of death coded using the Tenth International Classification of Diseases codes (ICD-10). The BMR included: age, sex, date of admission, date of death, HIV status, CD4 count, ART status and underlying cause of death (Table 1).

**Table 1:**
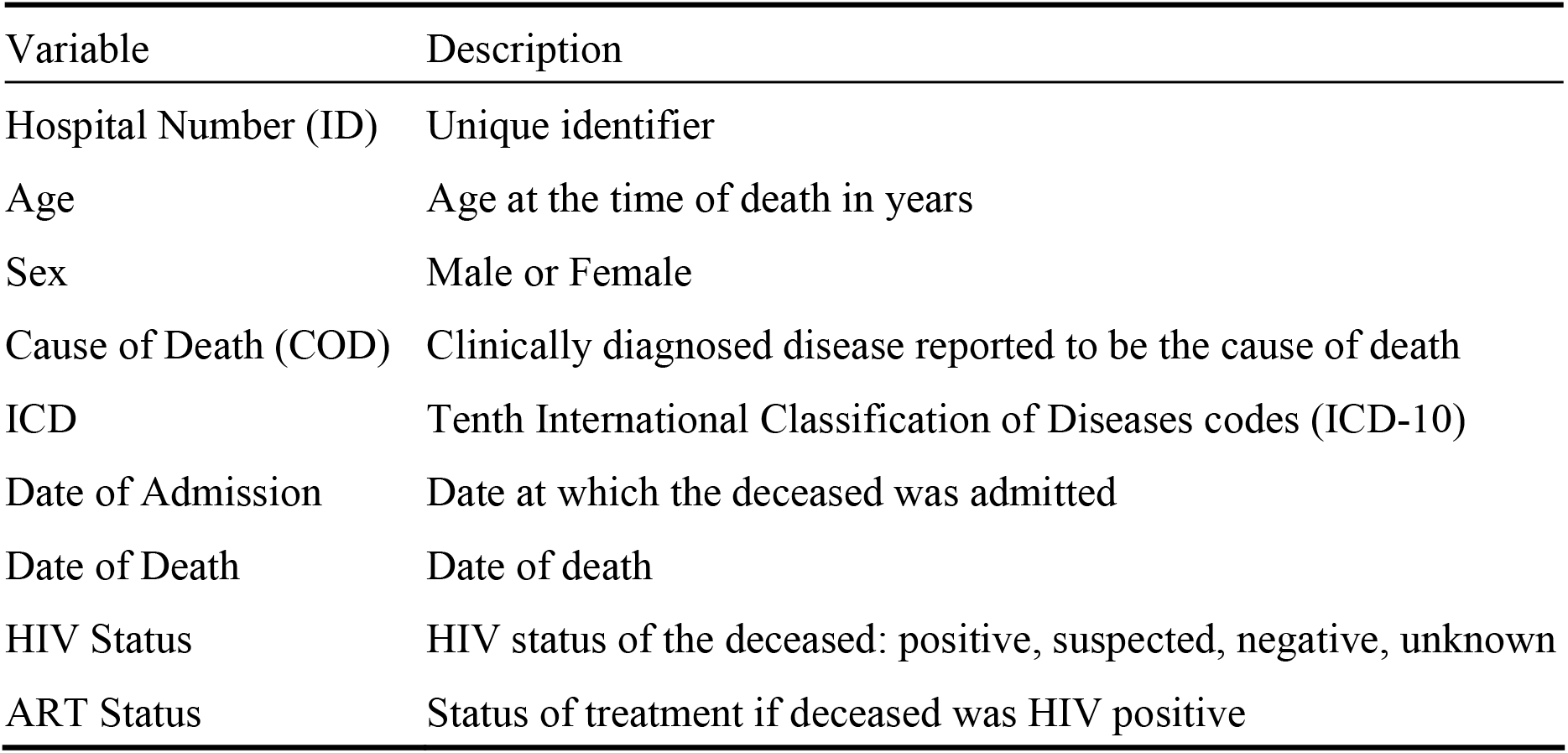
The main variables captured in the BMR

| Variable | Description |
| --- | --- |
| Hospital Number (ID) | Unique identifier |
| Age | Age at the time of death in years |
| Sex | Male or Female |
| Cause of Death (COD) | Clinically diagnosed disease reported to be the cause of death |
| ICD | Tenth International Classification of Diseases codes (ICD-10) |
| Date of Admission | Date at which the deceased was admitted |
| Date of Death | Date of death |
| HIV Status | HIV status of the deceased: positive, suspected, negative, unknown |
| ART Status | Status of treatment if deceased was HIV positive |

#### Duplicate Patient ID numbers

In the case of duplicate patient IDs, information in other fields was inspected. If the other fields suggested that they were different individuals, a new ID was assigned, if they appeared to be the same individual only one record was kept.

#### HIV and ART Status

A person’s HIV status was recorded as positive, negative, unknown, or clinically suspected because of an AIDS-defining illness. If the HIV status was not given but patients were on ART they were assumed to be HIV positive and if there was no information on ART status the HIV-status was given as ‘unknown’.

### Appendix 2: ICD codes in the data set

The ICD-10 codes in the data set are as follows:

A02-A07: Salmonella, shigellosis, amoebiasis, giardiasis
A09: Infectious gastroenteritis and colitis
A15-A16: Tuberculosis: extra-pulmonary
A17-A31: Tuberculosis: pulmonary
A32-A87: Infectious and parasitic diseases including those of the central nervous system
B01-B18: Infectious and parasitic diseases including those of the skin and viral hepatitis
B21-B24: HIV
B37-B46: Infectious and parasitic diseases: mycoses
B50-B59: Infectious and parasitic diseases: protozoal including mycoses
B69-B96: Infectious and parasitic diseases: helminthiasis and streptococcus
C02-C95: Malignant neoplasms
D00-D46: In-situ neoplasms
D51-D89: Diseases of the blood and blood-forming organs and certain disorders involving the immune mechanism
E03-E07: Disorders of thyroid gland
E10-E88: Certain disorders involving the immune mechanism including diabetes, endocrine glands and metabolic disorder
F01-F30: Mental and behavioural disorders
G00-G08: Inflammatory diseases of the nervous system including meningitis
G10-G99: Diseases of the nervous system
H52-H70: Diseases of the eyes and ears
I05-I95: Heart and cerebro-vascular diseases
J01-J99: Respiratory system
K25-K95: Digestive system
L03-L95: Skin and subcutaneous
M01-M99: Musculoskeletal
N00-N40: Genitourinary
O90-P20: Pregnancy
Q24-Z91: Congenital, injuries poison, external causes

#### Grouping the ICD codes

In order to establish control conditions, we excluded:

- Conditions for which having HIV is a risk factor for acquiring a disease which may be fatal.
- Conditions which share the same mode of transmission as HIV which may be fatal independent of HIV status but will have a higher prevalence in HIV infected persons and possibly a higher mortality.
- Diseases that may or may not be HIV related where HIV may cause the disease but there are multiple other causes that can not always be excluded.
- Diseases unrelated to HIV status but with a higher mortality in patients with HIV infection.

Code 0 refers to deaths that were omitted from the control group as there was uncertainty as to whether or not these causes of death were likely to be associated with HIV; code 1 refers to control conditions and codes 2 to 16 to conditions that are thought to be associated with HIV. Deaths that were not assigned an ICD code (529 or 3.4% of the sample) were excluded from the analysis. The codes were grouped as shown in Table 2.

**Table 2.**
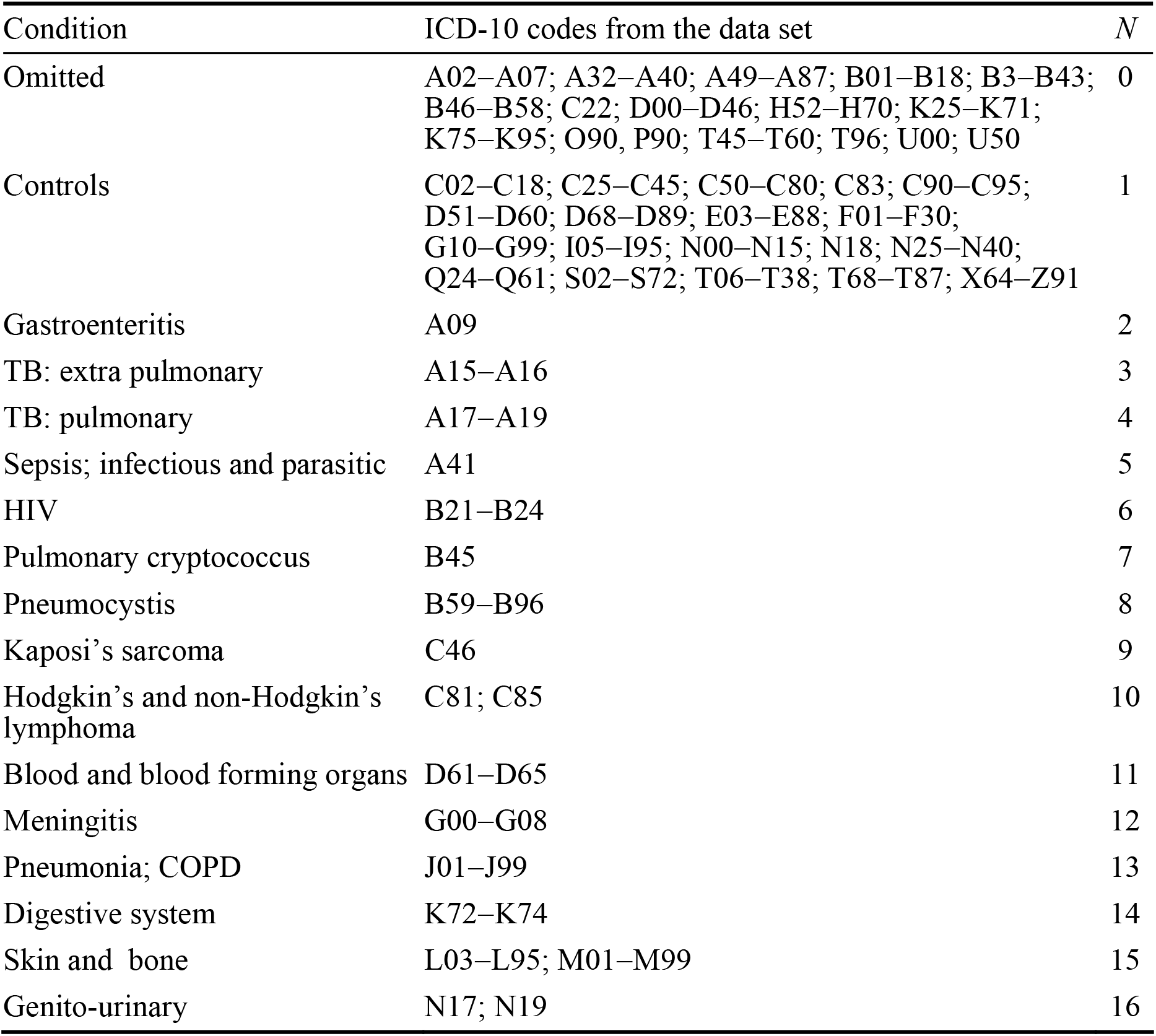
ICD-10 codes that were omitted, included in the control group, and included in disease categories. The right-most column is a code number corresponding to the ‘Assigned’ column in Table 3.

| Condition | ICD-10 codes from the data set | *N* |
| --- | --- | --- |
| Omitted | A02–A07; A32–A40; A49–A87; B01–B18; B3–B43; B46–B58; C22; D00–D46; H52–H70; K25–K71; K75–K95; O90, P90; T45–T60; T96; U00; U50 | 0 |
| Controls | C02–C18; C25–C45; C50–C80; C83; C90–C95; D51–D60; D68–D89; E03–E88; F01–F30; G10–G99; I05–I95; N00–N15; N18; N25–N40; Q24–Q61; S02–S72; T06–T38; T68–T87; X64–Z91 | 1 |
| Gastroenteritis | A09 | 2 |
| TB: extra pulmonary | A15–A16 | 3 |
| TB: pulmonary | A17–A19 | 4 |
| Sepsis; infectious and parasitic | A41 | 5 |
| HIV | B21–B24 | 6 |
| Pulmonary cryptococcus | B45 | 7 |
| Pneumocystis | B59–B96 | 8 |
| Kaposi’s sarcoma | C46 | 9 |
| Hodgkin’s and non-Hodgkin’s lymphoma | C81; C85 | 10 |
| Blood and blood forming organs | D61–D65 | 11 |
| Meningitis | G00–G08 | 12 |
| Pneumonia; COPD | J01–J99 | 13 |
| Digestive system | K72–K74 | 14 |
| Skin and bone | L03–L95; M01–M99 | 15 |
| Genito-urinary | N17; N19 | 16 |

Table 3 gives the data for each ICD-10 code and the last column gives the group to which each code was assigned as shown in Table 2. The grouped data are given in Table 4.

**Table 3.** Number of deaths, HIV-status, mean age, number of men and women, and the groups to which they are assigned as indicated in Table 2.

| ICD | Number | Negative | Positive | Unknown | Suspected | Mean age | Male | Female | Assigned |
| --- | --- | --- | --- | --- | --- | --- | --- | --- | --- |
| A02 | 24 | 1 | 21 | 1 | 1 | 39 | 14 | 0 | 0 |
| A03 | 1 | 0 | 1 | 0 | 0 | 32 | 0 | 1 | 0 |
| A06 | 8 | 1 | 5 | 2 | 0 | 52 | 3 | 5 | 0 |
| A07 | 1 | 0 | 1 | 0 | 0 | 39 | 0 | 1 | 0 |
| A09 | 509 | 19 | 366 | 61 | 63 | 43 | 220 | 289 | 2 |
| A15 | 685 | 18 | 565 | 48 | 54 | 39 | 367 | 318 | 3 |
| A16 | 736 | 38 | 557 | 64 | 77 | 41 | 394 | 342 | 3 |
| A17 | 364 | 10 | 295 | 25 | 34 | 38 | 197 | 318 | 4 |
| A18 | 473 | 17 | 394 | 23 | 39 | 39 | 246 | 342 | 4 |
| A19 | 95 | 4 | 82 | 3 | 6 | 39 | 49 | 46 | 4 |
| A31 | 17 | 0 | 16 | 0 | 1 | 39 | 10 | 7 | 4 |
| A32 | 1 | 1 | 0 | 0 | 0 | 28 | 1 | 0 | 0 |
| A35 | 2 | 0 | 0 | 2 | 0 | 42 | 2 | 0 | 0 |
| A39 | 3 | 0 | 2 | 1 | 0 | 31 | 1 | 2 | 0 |
| A40 | 4 | 0 | 1 | 2 | 1 | 54 | 3 | 1 | 0 |
| A41 | 889 | 152 | 454 | 194 | 89 | 49 | 386 | 503 | 5 |
| A49 | 2 | 0 | 2 | 0 | 0 | 41 | 1 | 1 | 0 |
| A52 | 1 | 0 | 0 | 1 | 0 | 32 | 1 | 0 | 0 |
| A81 | 6 | 0 | 6 | 0 | 0 | 36 | 3 | 3 | 0 |
| A85 | 1 | 0 | 1 | 0 | 0 | 32 | 0 | 1 | 0 |
| A86 | 1 | 0 | 0 | 0 | 1 | 36 | 1 | 0 | 0 |
| A87 | 1 | 0 | 1 | 0 | 0 | 36 | 0 | 1 | 0 |
| B01 | 20 | 1 | 11 | 4 | 4 | 30 | 4 | 16 | 0 |
| B02 | 1 | 0 | 1 | 0 | 0 | 34 | 0 | 1 | 0 |
| B10 | 7 | 0 | 4 | 2 | 1 | 47 | 2 | 5 | 0 |
| B16 | 9 | 2 | 5 | 2 | 0 | 42 | 4 | 5 | 0 |
| B17 | 2 | 0 | 1 | 1 | 0 | 47 | 1 | 1 | 0 |
| B18 | 1 | 1 | 0 | 0 | 0 | 57 | 1 | 0 | 0 |
| B21 | 4 | 0 | 4 | 0 | 0 | 31 | 2 | 2 | 6 |
| B22 | 6 | 2 | 3 | 0 | 1 | 42 | 3 | 3 | 6 |
| B23 | 7 | 0 | 7 | 0 | 0 | 35 | 4 | 3 | 6 |
| B24 | 656 | 5 | 601 | 5 | 45 | 39 | 319 | 337 | 6 |
| B37 | 2 | 0 | 1 | 1 | 0 | 70 | 0 | 2 | 0 |
| B39 | 2 | 0 | 2 | 0 | 0 | 31 | 2 | 0 | 0 |
| B42 | 1 | 0 | 1 | 0 | 0 | 28 | 0 | 1 | 0 |
| B43 | 17 | 5 | 9 | 2 | 1 | 40 | 11 | 6 | 0 |
| B45 | 353 | 1 | 323 | 6 | 23 | 37 | 168 | 185 | 7 |
| B46 | 1 | 0 | 0 | 1 | 0 | 57 | 0 | 1 | 0 |
| B50 | 3 | 0 | 0 | 2 | 1 | 39 | 2 | 1 | 0 |
| B54 | 9 | 2 | 2 | 3 | 2 | 41 | 4 | 5 | 0 |
| B58 | 5 | 0 | 5 | 0 | 0 | 39 | 4 | 1 | 0 |
| B59 | 170 | 2 | 149 | 6 | 13 | 37 | 60 | 110 | 8 |
| B69 | 1 | 0 | 1 | 0 | 0 | 61 | 0 | 1 | 8 |
| B95 | 6 | 0 | 6 | 0 | 0 | 48 | 3 | 3 | 8 |
| B96 | 43 | 6 | 28 | 6 | 3 | 48 | 21 | 22 | 8 |
| C02 | 1 | 0 | 0 | 1 | 0 | 59 | 1 | 0 | 1 |
| C04 | 1 | 0 | 1 | 0 | 0 | 48 | 0 | 1 | 1 |
| C10 | 1 | 0 | 0 | 1 | 0 | 45 | 1 | 0 | 1 |
| C15 | 4 | 1 | 0 | 2 | 1 | 73 | 4 | 0 | 1 |
| C16 | 1 | 0 | 0 | 1 | 0 | 67 | 1 | 0 | 1 |
| C18 | 1 | 0 | 1 | 0 | 0 | 55 | 1 | 0 | 1 |
| C22 | 34 | 9 | 5 | 14 | 6 | 60 | 20 | 14 | 0 |
| C25 | 15 | 3 | 0 | 10 | 2 | 62 | 7 | 8 | 1 |
| C26 | 1 | 0 | 0 | 1 | 0 | 69 | 1 | 0 | 1 |
| C32 | 2 | 0 | 0 | 2 | 0 | 66 | 2 | 0 | 1 |
| C34 | 151 | 86 | 9 | 45 | 11 | 61 | 121 | 30 | 1 |
| C45 | 2 | 1 | 0 | 1 | 0 | 65 | 1 | 1 | 1 |
| C46 | 117 | 3 | 108 | 2 | 4 | 37 | 63 | 54 | 9 |
| C50 | 34 | 9 | 7 | 13 | 5 | 59 | 0 | 34 | 1 |
| C56 | 6 | 1 | 1 | 3 | 1 | 58 | 0 | 6 | 1 |
| C61 | 58 | 26 | 0 | 24 | 8 | 72 | 58 | 0 | 1 |
| C62 | 2 | 1 | 0 | 1 | 0 | 35 | 2 | 0 | 1 |
| C64 | 4 | 2 | 1 | 1 | 0 | 52 | 3 | 1 | 1 |
| C67 | 7 | 2 | 0 | 3 | 2 | 63 | 7 | 0 | 1 |
| C70 | 4 | 0 | 0 | 4 | 0 | 61 | 1 | 3 | 1 |
| C71 | 3 | 1 | 0 | 2 | 0 | 52 | 1 | 2 | 1 |
| C72 | 1 | 0 | 1 | 0 | 0 | 31 | 0 | 1 | 1 |
| C74 | 1 | 0 | 1 | 0 | 0 | 32 | 1 | 0 | 1 |
| C76 | 2 | 0 | 0 | 2 | 0 | 68 | 2 | 0 | 1 |
| C78 | 1 | 1 | 0 | 0 | 0 | 73 | 1 | 0 | 1 |
| C79 | 25 | 10 | 0 | 10 | 5 | 64 | 17 | 8 | 1 |
| C80 | 95 | 39 | 7 | 40 | 9 | 65 | 42 | 53 | 1 |
| C81 | 41 | 12 | 25 | 4 | 0 | 40 | 19 | 22 | 10 |
| C83 | 2 | 1 | 0 | 1 | 0 | 57 | 2 | 0 | 1 |
| C85 | 54 | 5 | 44 | 3 | 2 | 40 | 28 | 26 | 10 |
| C90 | 40 | 20 | 3 | 12 | 5 | 61 | 18 | 22 | 1 |
| C91 | 17 | 8 | 3 | 6 | 0 | 59 | 8 | 9 | 1 |
| C92 | 29 | 12 | 3 | 11 | 3 | 49 | 12 | 17 | 1 |
| C94 | 3 | 1 | 1 | 1 | 0 | 67 | 2 | 1 | 1 |
| C95 | 12 | 6 | 2 | 3 | 1 | 40 | 9 | 3 | 1 |
| D00 | 19 | 8 | 1 | 4 | 6 | 62 | 13 | 6 | 0 |
| D01 | 9 | 1 | 4 | 4 | 0 | 59 | 5 | 4 | 0 |
| D02 | 1 | 0 | 0 | 0 | 1 | 94 | 1 | 0 | 0 |
| D03 | 2 | 1 | 0 | 1 | 0 | 67 | 1 | 1 | 0 |
| D06 | 29 | 2 | 11 | 12 | 4 | 57 | 0 | 29 | 0 |
| D07 | 6 | 1 | 1 | 2 | 2 | 68 | 0 | 6 | 0 |
| D09 | 1 | 0 | 0 | 1 | 0 | 83 | 1 | 0 | 0 |
| D15 | 1 | 1 | 0 | 0 | 0 | 20 | 0 | 1 | 0 |
| D43 | 2 | 0 | 1 | 1 | 0 | 49 | 0 | 2 | 0 |
| D46 | 1 | 1 | 0 | 0 | 0 | 85 | 0 | 1 | 0 |
| D51 | 2 | 2 | 0 | 0 | 0 | 51 | 0 | 2 | 1 |
| D53 | 1 | 0 | 0 | 1 | 0 | 63 | 0 | 1 | 1 |
| D59 | 1 | 1 | 0 | 0 | 0 | 28 | 0 | 1 | 1 |
| D60 | 2 | 0 | 2 | 0 | 0 | 39 | 0 | 2 | 1 |
| D61 | 37 | 4 | 31 | 0 | 2 | 36 | 24 | 13 | 11 |
| D64 | 85 | 6 | 51 | 20 | 8 | 43 | 32 | 53 | 11 |
| D65 | 14 | 2 | 10 | 2 | 0 | 37 | 5 | 9 | 11 |
| D68 | 20 | 13 | 1 | 5 | 1 | 38 | 4 | 16 | 1 |
| D69 | 15 | 1 | 12 | 1 | 1 | 41 | 5 | 10 | 1 |
| D70 | 3 | 0 | 1 | 2 | 0 | 54 | 1 | 2 | 1 |
| D73 | 1 | 0 | 1 | 0 | 0 | 40 | 0 | 1 | 1 |
| D75 | 1 | 0 | 1 | 0 | 0 | 51 | 0 | 1 | 1 |
| D81 | 1 | 0 | 0 | 0 | 1 | 65 | 1 | 0 | 1 |
| D89 | 6 | 0 | 5 | 0 | 1 | 35 | 0 | 6 | 1 |
| E03 | 5 | 2 | 0 | 3 | 0 | 78 | 2 | 3 | 1 |
| E05 | 3 | 0 | 0 | 3 | 0 | 52 | 1 | 2 | 1 |
| E07 | 1 | 1 | 0 | 0 | 0 | 49 | 1 | 0 | 1 |
| E10 | 3 | 0 | 0 | 2 | 1 | 75 | 0 | 3 | 1 |
| E11 | 11 | 3 | 1 | 7 | 0 | 63 | 3 | 8 | 1 |
| E12 | 24 | 8 | 1 | 10 | 5 | 68 | 11 | 13 | 1 |
| E14 | 164 | 52 | 14 | 73 | 25 | 60 | 64 | 100 | 1 |
| E15 | 5 | 1 | 0 | 4 | 0 | 70 | 2 | 3 | 1 |
| E16 | 134 | 25 | 29 | 59 | 21 | 58 | 67 | 67 | 1 |
| E20 | 3 | 0 | 0 | 3 | 0 | 64 | 3 | 0 | 1 |
| E21 | 7 | 3 | 1 | 3 | 0 | 66 | 5 | 2 | 1 |
| E22 | 2 | 0 | 1 | 0 | 1 | 39 | 2 | 0 | 1 |
| E23 | 1 | 0 | 0 | 1 | 0 | 54 | 1 | 0 | 1 |
| E27 | 4 | 0 | 4 | 0 | 0 | 38 | 2 | 2 | 1 |
| E46 | 2 | 0 | 0 | 2 | 0 | 57 | 1 | 1 | 1 |
| E52 | 1 | 0 | 0 | 1 | 0 | 38 | 0 | 1 | 1 |
| E66 | 10 | 4 | 1 | 4 | 1 | 59 | 2 | 8 | 1 |
| E77 | 3 | 1 | 0 | 1 | 1 | 62 | 1 | 2 | 1 |
| E84 | 1 | 0 | 1 | 0 | 0 | 33 | 1 | 0 | 1 |
| E86 | 81 | 8 | 37 | 18 | 18 | 51 | 48 | 33 | 1 |
| E87 | 142 | 5 | 117 | 12 | 8 | 41 | 34 | 108 | 1 |
| E88 | 6 | 2 | 2 | 1 | 1 | 63 | 3 | 3 | 1 |
| F01 | 10 | 4 | 1 | 4 | 1 | 73 | 6 | 4 | 1 |
| F02 | 8 | 0 | 7 | 1 | 0 | 47 | 3 | 5 | 1 |
| F03 | 15 | 4 | 5 | 6 | 0 | 63 | 7 | 8 | 1 |
| F05 | 12 | 0 | 4 | 5 | 3 | 57 | 7 | 5 | 1 |
| F07 | 1 | 1 | 0 | 0 | 0 | 66 | 0 | 1 | 1 |
| F10 | 1 | 0 | 1 | 0 | 0 | 49 | 1 | 0 | 1 |
| F30 | 1 | 0 | 1 | 0 | 0 | 29 | 0 | 1 | 1 |
| G00 | 79 | 5 | 46 | 17 | 11 | 40 | 31 | 48 | 12 |
| G03 | 593 | 24 | 388 | 109 | 72 | 40 | 294 | 299 | 12 |
| G04 | 5 | 0 | 5 | 0 | 0 | 46 | 3 | 2 | 12 |
| G06 | 5 | 1 | 2 | 1 | 1 | 48 | 2 | 3 | 12 |
| G08 | 2 | 0 | 1 | 1 | 0 | 54 | 0 | 2 | 12 |
| G10 | 1 | 1 | 0 | 0 | 0 | 51 | 0 | 1 | 1 |
| G11 | 2 | 0 | 0 | 2 | 0 | 50 | 2 | 0 | 1 |
| G12 | 2 | 1 | 0 | 1 | 0 | 52 | 2 | 0 | 1 |
| G20 | 3 | 2 | 0 | 1 | 0 | 76 | 3 | 0 | 1 |
| G21 | 3 | 0 | 1 | 2 | 0 | 29 | 2 | 1 | 1 |
| G23 | 1 | 0 | 0 | 1 | 0 | 90 | 0 | 1 | 1 |
| G30 | 2 | 1 | 0 | 1 | 0 | 75 | 1 | 1 | 1 |
| G31 | 1 | 0 | 0 | 1 | 0 | 54 | 1 | 0 | 1 |
| G37 | 10 | 0 | 6 | 4 | 0 | 44 | 5 | 5 | 1 |
| G40 | 55 | 10 | 13 | 21 | 11 | 49 | 38 | 17 | 1 |
| G41 | 32 | 7 | 4 | 18 | 3 | 50 | 17 | 15 | 1 |
| G45 | 2 | 1 | 0 | 1 | 0 | 68 | 1 | 1 | 1 |
| G47 | 2 | 1 | 0 | 1 | 0 | 72 | 1 | 1 | 1 |
| G58 | 1 | 0 | 0 | 0 | 1 | 54 | 1 | 0 | 1 |
| G61 | 7 | 0 | 4 | 3 | 0 | 43 | 3 | 4 | 1 |
| G70 | 1 | 1 | 0 | 0 | 0 | 35 | 0 | 1 | 1 |
| G71 | 1 | 0 | 0 | 1 | 0 | 15 | 1 | 0 | 1 |
| G72 | 1 | 0 | 1 | 0 | 0 | 27 | 1 | 0 | 1 |
| G81 | 12 | 3 | 3 | 3 | 3 | 60 | 5 | 7 | 1 |
| G82 | 6 | 1 | 1 | 3 | 1 | 60 | 2 | 4 | 1 |
| G83 | 1 | 1 | 0 | 0 | 0 | 82 | 0 | 1 | 1 |
| G90 | 2 | 0 | 2 | 0 | 0 | 62 | 0 | 2 | 1 |
| G91 | 16 | 2 | 9 | 2 | 3 | 42 | 8 | 8 | 1 |
| G93 | 26 | 4 | 9 | 12 | 1 | 46 | 13 | 13 | 1 |
| G94 | 1 | 1 | 0 | 0 | 0 | 51 | 1 | 0 | 1 |
| G95 | 3 | 0 | 3 | 0 | 0 | 40 | 1 | 2 | 1 |
| G99 | 2 | 1 | 1 | 0 | 0 | 34 | 0 | 2 | 1 |
| H52 | 1 | 0 | 1 | 0 | 0 | 34 | 1 | 0 | 0 |
| H66 | 1 | 0 | 1 | 0 | 0 | 44 | 0 | 1 | 0 |
| H70 | 1 | 0 | 1 | 0 | 0 | 47 | 0 | 1 | 0 |
| I05 | 13 | 6 | 1 | 6 | 0 | 55 | 2 | 11 | 1 |
| I06 | 2 | 2 | 0 | 0 | 0 | 51 | 1 | 1 | 1 |
| i08 | 6 | 3 | 0 | 3 | 0 | 40 | 3 | 3 | 1 |
| I09 | 8 | 5 | 1 | 1 | 1 | 51 | 4 | 4 | 1 |
| I10 | 106 | 25 | 6 | 60 | 15 | 65 | 33 | 73 | 1 |
| I11 | 327 | 123 | 25 | 138 | 41 | 65 | 113 | 214 | 1 |
| I21 | 40 | 11 | 3 | 20 | 6 | 63 | 22 | 18 | 1 |
| I22 | 2 | 1 | 0 | 1 | 0 | 70 | 2 | 0 | 1 |
| I24 | 5 | 2 | 0 | 3 | 0 | 71 | 1 | 4 | 1 |
| I25 | 12 | 4 | 0 | 6 | 2 | 67 | 6 | 6 | 1 |
| I26 | 115 | 30 | 37 | 39 | 9 | 51 | 35 | 80 | 1 |
| I27 | 198 | 62 | 37 | 84 | 15 | 59 | 96 | 102 | 1 |
| I30 | 3 | 1 | 2 | 0 | 0 | 35 | 2 | 1 | 1 |
| I31 | 15 | 3 | 5 | 3 | 4 | 40 | 8 | 7 | 1 |
| I33 | 2 | 1 | 1 | 0 | 0 | 25 | 1 | 1 | 1 |
| I35 | 1 | 0 | 0 | 0 | 1 | 51 | 0 | 1 | 1 |
| I37 | 2 | 2 | 0 | 0 | 0 | 41 | 0 | 2 | 1 |
| I38 | 12 | 3 | 4 | 3 | 2 | 45 | 6 | 6 | 1 |
| I42 | 231 | 58 | 60 | 106 | 7 | 53 | 116 | 115 | 1 |
| I44 | 7 | 1 | 0 | 5 | 1 | 65 | 2 | 5 | 1 |
| I46 | 3 | 2 | 0 | 1 | 0 | 93 | 0 | 3 | 1 |
| I47 | 5 | 2 | 0 | 3 | 0 | 65 | 3 | 2 | 1 |
| I48 | 14 | 4 | 1 | 6 | 3 | 62 | 4 | 10 | 1 |
| I49 | 1 | 1 | 0 | 0 | 0 | 68 | 0 | 1 | 1 |
| I50 | 318 | 105 | 36 | 143 | 34 | 62 | 123 | 195 | 1 |
| I51 | 5 | 0 | 1 | 4 | 0 | 49 | 3 | 2 | 1 |
| I60 | 1 | 0 | 0 | 1 | 0 | 51 | 0 | 1 | 1 |
| I61 | 202 | 48 | 20 | 119 | 15 | 57 | 97 | 105 | 1 |
| I62 | 25 | 6 | 3 | 16 | 0 | 57 | 16 | 9 | 1 |
| I63 | 38 | 7 | 11 | 15 | 5 | 62 | 16 | 22 | 1 |
| I64 | 998 | 331 | 108 | 461 | 98 | 64 | 390 | 608 | 1 |
| I71 | 4 | 1 | 1 | 2 | 0 | 53 | 1 | 3 | 1 |
| I72 | 1 | 0 | 0 | 1 | 0 | 87 | 1 | 0 | 1 |
| I73 | 7 | 2 | 1 | 4 | 0 | 63 | 2 | 5 | 1 |
| I80 | 22 | 2 | 13 | 6 | 1 | 48 | 10 | 12 | 1 |
| I82 | 3 | 1 | 2 | 0 | 0 | 30 | 1 | 2 | 1 |
| I85 | 2 | 1 | 0 | 1 | 0 | 66 | 0 | 2 | 1 |
| I88 | 1 | 0 | 0 | 0 | 1 | 25 | 0 | 1 | 1 |
| I95 | 3 | 1 | 1 | 1 | 0 | 50 | 2 | 1 | 1 |
| J01 | 1 | 0 | 1 | 0 | 0 | 31 | 0 | 1 | 13 |
| J06 | 2 | 0 | 2 | 0 | 0 | 29 | 0 | 2 | 13 |
| J13 | 4 | 1 | 3 | 0 | 0 | 45 | 2 | 2 | 13 |
| J15 | 4 | 0 | 2 | 1 | 1 | 53 | 3 | 1 | 13 |
| J17 | 1 | 0 | 1 | 0 | 0 | 29 | 0 | 1 | 13 |
| J18 | 2198 | 210 | 1206 | 453 | 329 | 46 | 1047 | 1151 | 13 |
| J22 | 233 | 11 | 140 | 38 | 44 | 45 | 126 | 107 | 13 |
| J35 | 2 | 1 | 1 | 0 | 0 | 44 | 0 | 2 | 13 |
| J43 | 2 | 1 | 1 | 0 | 0 | 58 | 1 | 1 | 13 |
| J44 | 126 | 29 | 5 | 76 | 16 | 63 | 98 | 28 | 13 |
| J45 | 2 | 0 | 1 | 0 | 1 | 52 | 1 | 1 | 13 |
| J46 | 23 | 6 | 4 | 9 | 4 | 57 | 11 | 12 | 13 |
| J47 | 22 | 10 | 8 | 4 | 0 | 47 | 13 | 9 | 13 |
| J69 | 40 | 6 | 5 | 25 | 4 | 52 | 27 | 13 | 13 |
| J81 | 19 | 5 | 1 | 9 | 4 | 56 | 9 | 10 | 13 |
| J84 | 5 | 3 | 1 | 0 | 1 | 63 | 2 | 3 | 13 |
| J85 | 15 | 4 | 7 | 3 | 1 | 45 | 11 | 4 | 13 |
| J86 | 37 | 5 | 29 | 3 | 0 | 39 | 20 | 17 | 13 |
| J90 | 62 | 16 | 24 | 12 | 10 | 49 | 33 | 29 | 13 |
| J91 | 2 | 0 | 1 | 1 | 0 | 36 | 1 | 1 | 13 |
| J93 | 20 | 2 | 13 | 3 | 2 | 47 | 16 | 4 | 13 |
| J94 | 2 | 0 | 2 | 0 | 0 | 39 | 0 | 2 | 13 |
| J96 | 31 | 10 | 9 | 6 | 6 | 53 | 20 | 11 | 13 |
| J98 | 3 | 1 | 1 | 0 | 1 | 62 | 2 | 1 | 13 |
| J99 | 2 | 0 | 1 | 1 | 0 | 50 | 1 | 1 | 13 |
| K25 | 4 | 1 | 0 | 3 | 0 | 48 | 4 | 0 | 0 |
| K27 | 6 | 3 | 0 | 3 | 0 | 68 | 2 | 4 | 0 |
| K29 | 3 | 0 | 2 | 1 | 0 | 55 | 2 | 1 | 0 |
| K31 | 2 | 0 | 1 | 1 | 0 | 28 | 1 | 1 | 0 |
| K42 | 1 | 1 | 0 | 0 | 0 | 80 | 0 | 1 | 0 |
| K56 | 9 | 1 | 1 | 5 | 2 | 55 | 3 | 6 | 0 |
| K59 | 13 | 2 | 10 | 1 | 0 | 45 | 5 | 8 | 0 |
| K63 | 1 | 0 | 1 | 0 | 0 | 20 | 0 | 1 | 0 |
| K65 | 13 | 4 | 3 | 6 | 0 | 47 | 7 | 6 | 0 |
| K70 | 24 | 6 | 0 | 15 | 3 | 55 | 13 | 11 | 0 |
| K71 | 28 | 1 | 24 | 2 | 1 | 35 | 16 | 12 | 0 |
| K72 | 165 | 23 | 63 | 62 | 17 | 49 | 96 | 69 | 14 |
| K73 | 34 | 0 | 21 | 9 | 4 | 38 | 18 | 16 | 14 |
| K74 | 41 | 12 | 9 | 19 | 1 | 51 | 27 | 14 | 14 |
| K75 | 2 | 2 | 0 | 0 | 0 | 39 | 0 | 2 | 0 |
| K76 | 18 | 3 | 6 | 7 | 2 | 48 | 15 | 3 | 0 |
| K77 | 1 | 0 | 1 | 0 | 0 | 40 | 1 | 0 | 0 |
| K83 | 1 | 0 | 0 | 1 | 0 | 81 | 0 | 1 | 0 |
| K85 | 7 | 0 | 4 | 3 | 0 | 49 | 3 | 4 | 0 |
| K86 | 1 | 1 | 0 | 0 | 0 | 47 | 1 | 0 | 0 |
| K92 | 67 | 18 | 12 | 31 | 6 | 59 | 34 | 33 | 0 |
| K95 | 1 | 0 | 1 | 0 | 0 | 33 | 0 | 1 | 0 |
| L03 | 4 | 1 | 1 | 2 | 0 | 55 | 1 | 3 | 15 |
| L12 | 2 | 2 | 0 | 0 | 0 | 66 | 1 | 1 | 15 |
| L13 | 4 | 0 | 1 | 3 | 0 | 53 | 3 | 1 | 15 |
| L30 | 1 | 0 | 1 | 0 | 0 | 48 | 0 | 1 | 15 |
| L51 | 29 | 1 | 21 | 3 | 4 | 45 | 18 | 11 | 15 |
| L53 | 1 | 0 | 1 | 0 | 0 | 45 | 0 | 1 | 15 |
| L56 | 1 | 0 | 1 | 0 | 0 | 35 | 0 | 1 | 15 |
| L89 | 3 | 1 | 0 | 2 | 0 | 54 | 1 | 2 | 15 |
| L95 | 1 | 0 | 0 | 1 | 0 | 39 | 0 | 1 | 15 |
| M01 | 1 | 0 | 1 | 0 | 0 | 38 | 1 | 0 | 15 |
| M07 | 1 | 0 | 0 | 1 | 0 | 63 | 0 | 1 | 15 |
| M19 | 1 | 1 | 0 | 0 | 0 | 57 | 0 | 1 | 15 |
| M31 | 14 | 3 | 8 | 1 | 2 | 35 | 3 | 11 | 15 |
| M33 | 2 | 1 | 0 | 1 | 0 | 60 | 1 | 1 | 15 |
| M34 | 3 | 1 | 0 | 1 | 1 | 32 | 0 | 3 | 15 |
| M35 | 1 | 0 | 1 | 0 | 0 | 33 | 0 | 1 | 15 |
| M49 | 1 | 0 | 1 | 0 | 0 | 22 | 0 | 1 | 15 |
| M72 | 2 | 1 | 0 | 1 | 0 | 49 | 1 | 1 | 15 |
| M80 | 12 | 4 | 4 | 4 | 0 | 53 | 8 | 4 | 15 |
| M81 | 54 | 21 | 8 | 19 | 6 | 58 | 36 | 18 | 15 |
| M83 | 1 | 1 | 0 | 0 | 0 | 47 | 1 | 0 | 15 |
| M85 | 1 | 0 | 0 | 0 | 1 | 45 | 1 | 0 | 15 |
| M86 | 1 | 0 | 1 | 0 | 0 | 46 | 0 | 1 | 15 |
| M89 | 1 | 0 | 0 | 0 | 1 | 22 | 1 | 0 | 15 |
| M98 | 3 | 1 | 1 | 1 | 0 | 41 | 3 | 0 | 15 |
| M99 | 2 | 0 | 2 | 0 | 0 | 35 | 1 | 1 | 15 |
| N00 | 3 | 1 | 1 | 1 | 0 | 37 | 2 | 1 | 1 |
| N04 | 6 | 2 | 3 | 1 | 0 | 27 | 1 | 5 | 1 |
| N05 | 1 | 0 | 0 | 1 | 0 | 58 | 0 | 1 | 1 |
| N11 | 5 | 2 | 1 | 1 | 1 | 44 | 2 | 3 | 1 |
| N13 | 11 | 5 | 1 | 5 | 0 | 76 | 9 | 2 | 1 |
| N14 | 2 | 1 | 0 | 0 | 1 | 57 | 2 | 0 | 1 |
| N15 | 1 | 0 | 1 | 0 | 0 | 61 | 1 | 0 | 1 |
| N17 | 153 | 28 | 83 | 28 | 14 | 49 | 77 | 76 | 16 |
| N18 | 13 | 6 | 3 | 2 | 2 | 50 | 7 | 6 | 1 |
| N19 | 722 | 161 | 295 | 193 | 73 | 51 | 384 | 338 | 16 |
| N25 | 2 | 2 | 0 | 0 | 0 | 61 | 0 | 2 | 1 |
| N39 | 10 | 3 | 0 | 4 | 3 | 76 | 3 | 7 | 1 |
| NA0 | 45 | 6 | 17 | 19 | 3 | 49 | 25 | 20 | 1 |
| O90 | 17 | 4 | 7 | 4 | 2 | 29 | 0 | 17 | 0 |
| P20 | 1 | 0 | 0 | 0 | 1 | 29 | 0 | 1 | 0 |
| Q24 | 1 | 0 | 1 | 0 | 0 | 24 | 0 | 1 | 1 |
| Q26 | 4 | 2 | 0 | 2 | 0 | 56 | 4 | 0 | 1 |
| Q61 | 1 | 1 | 0 | 0 | 0 | 56 | 1 | 0 | 1 |
| S02 | 1 | 0 | 0 | 1 | 0 | 14 | 1 | 0 | 1 |
| S06 | 72 | 19 | 4 | 45 | 4 | 53 | 27 | 45 | 1 |
| S72 | 1 | 0 | 0 | 1 | 0 | 63 | 1 | 0 | 1 |
| T06 | 1 | 0 | 0 | 0 | 1 | 35 | 1 | 0 | 1 |
| T38 | 1 | 0 | 0 | 1 | 0 | 38 | 0 | 1 | 1 |
| T45 | 8 | 2 | 0 | 6 | 0 | 59 | 3 | 5 | 0 |
| T48 | 1 | 0 | 0 | 1 | 0 | 21 | 0 | 1 | 0 |
| T59 | 1 | 0 | 0 | 1 | 0 | 67 | 1 | 0 | 0 |
| T60 | 27 | 1 | 1 | 20 | 5 | 33 | 17 | 10 | 0 |
| T68 | 11 | 2 | 1 | 7 | 1 | 63 | 4 | 7 | 1 |
| T78 | 4 | 1 | 1 | 2 | 0 | 56 | 0 | 4 | 1 |
| T87 | 1 | 0 | 0 | 1 | 0 | 76 | 0 | 1 | 1 |
| T96 | 4 | 1 | 1 | 2 | 0 | 44 | 2 | 2 | 0 |
| U00 | 1 | 0 | 0 | 0 | 1 | 70 | 0 | 1 | 0 |
| U50 | 21 | 1 | 16 | 1 | 3 | 38 | 10 | 11 | 0 |
| X64 | 3 | 0 | 0 | 3 | 0 | 29 | 2 | 1 | 1 |
| X66 | 1 | 0 | 1 | 0 | 0 | 26 | 0 | 1 | 1 |
| X69 | 1 | 0 | 0 | 1 | 0 | 34 | 0 | 1 | 1 |
| Y84 | 1 | 0 | 0 | 0 | 1 | 54 | 1 | 0 | 1 |
| Y91 | 9 | 2 | 0 | 7 | 0 | 59 | 6 | 3 | 1 |
| Y95 | 1 | 0 | 0 | 1 | 0 | 79 | 1 | 0 | 1 |
| Z03 | 91 | 3 | 71 | 12 | 5 | 41 | 50 | 41 | 1 |
| Z49 | 1 | 0 | 0 | 1 | 0 | 48 | 1 | 0 | 1 |
| Z91 | 1 | 0 | 0 | 1 | 0 | 54 | 0 | 1 | 1 |
| UC | 652 | 104 | 184 | 167 | 74 | 49 | 310 | 219 | -1 |

**Table 4.**
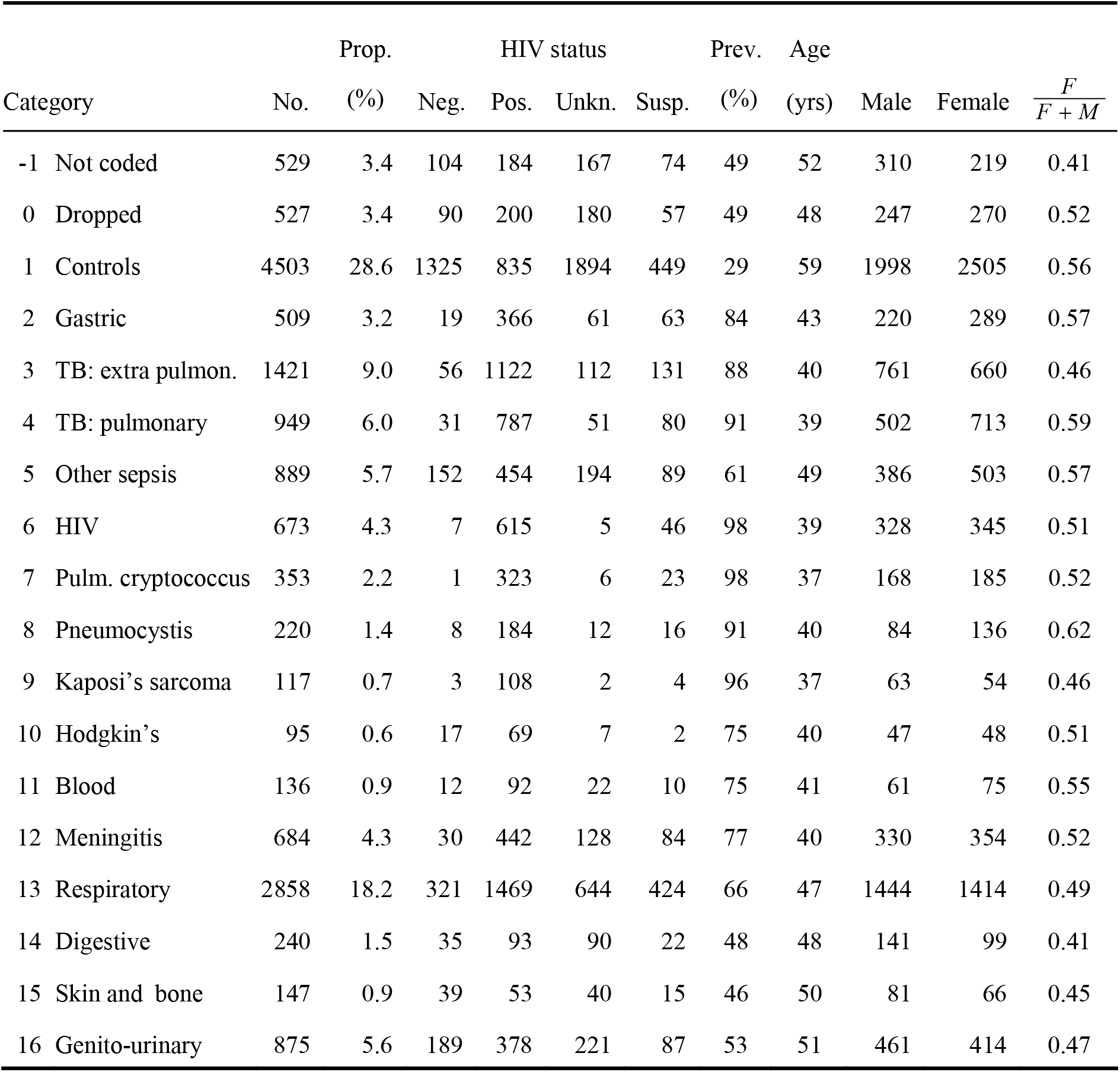
The data as grouped in Table 3. The columns give the number of deaths in each category, the proportion of all deaths attributable to each category, the number in each HIV-category (negative, positive, unknown or suspected), the prevalence of HIV assuming that all that are HIV-unknown are uninfected and all that are suspected of being infected with HIV are infected, the mean age, the number of men and women and the sex ratio.

### Appendix 3: Distribution of cases by age, sex and HIV-status

Each person that died was recorded as being HIV positive *(P)*, HIV-negative *(N)*, HIV suspected *(S)* or HIV unknown *(U)*. For those that were positive or negative we fitted the distribution of cases by age and gender to skew-normal distributions so that *P(a)* the number of people of HIV-positive people of age *a*, for each gender is given by

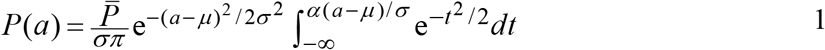

where 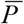 is the total number of people recorded as being HIV-positive, *μ* is the location parameter, *σ* is the scale parameter, and *α* determines the skewness of the distribution. A similar functional form is fitted to those that are recorded as being HIV-negative, *N(a)*. The data and the fitted curves are plotted in Figure 1.

**Figure 1.**
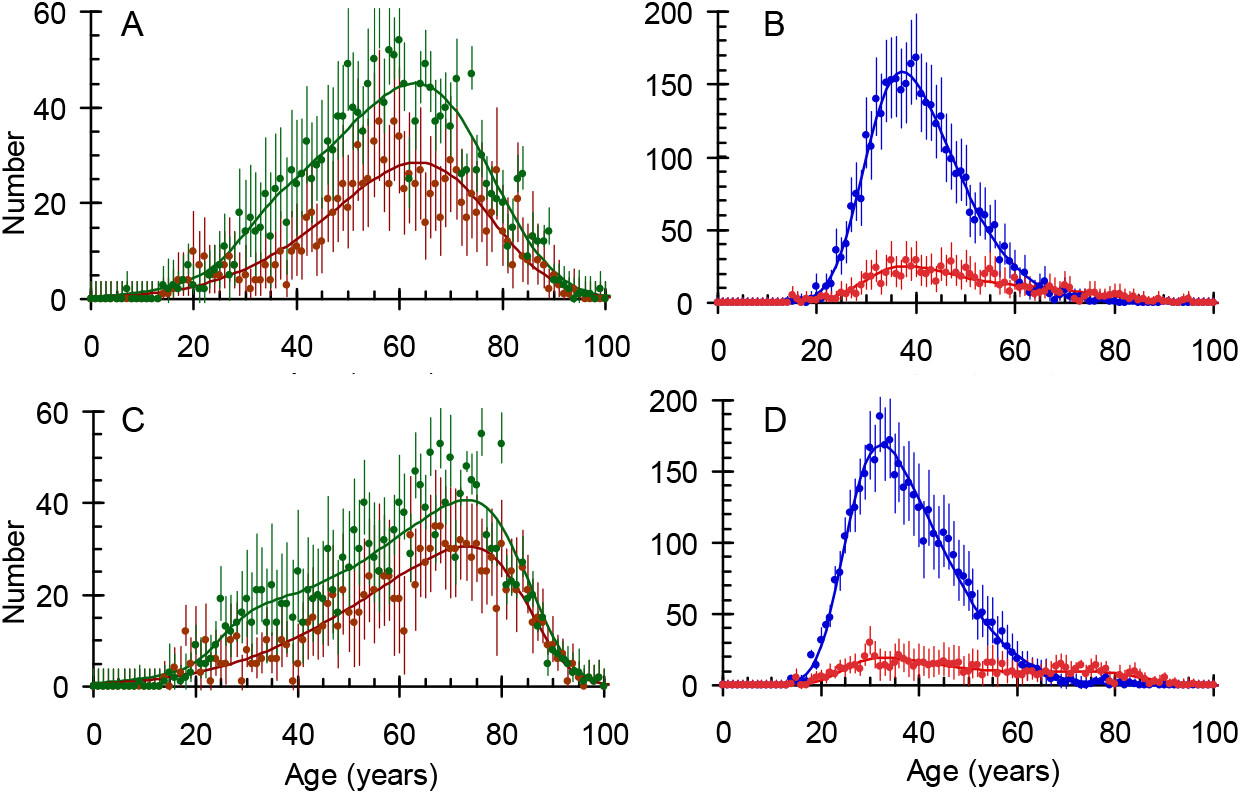
A and B: men; C and D women. Brown: HIV-negative; green: HIV unknown. Blue: HIV-positive; red: HIV-suspected.

The data in Figure 1 allow us to estimate the proportion of men and women who are HIV-positive and HIV-negative among those whose status is suspected or unknown by fitting a weighted sum of the fitted curves for those whose status is known to be positive or negative to those whose status is unknown or suspected. We then have

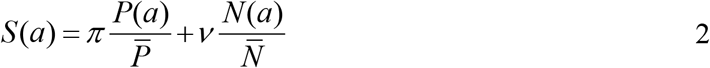

For those suspected of being HIV-positive *π* is the number that are HIV-positive, *ν* is the number that are HIV negative. A similar expression for *U*(*a*) gives the corresponding proportions for those whose HIV-status is unknown and we do this separately for men and women. The data and the fitted curves are given in Figure 1 and Table 5.

**Table 5.** Parameters for the skew-normal fits to the number of men (M) and women (W) who are HIV positive (P) or HIV negative (N).

|  | MP | MN | WP | WN |
| --- | --- | --- | --- | --- |
| Shape | 2.69 | -2.16 | 3.51 | -4.25 |
| Norm | 3672 | 1155 | 4102 | 1285 |
| Location | 29.9 | 76.4 | 24.9 | 85.1 |
| Scale | 14.9 | 24.5 | 16.5 | 29.6 |

The fits to the data in Figure 1 show that the distribution of HIV-negative men and women (brown lines) are both skewed to the left with the modal age at death for men being 64 years and for women 74 years while the distribution of HIV-positive men and women (blue lines) are both skew to the right with the modal age at death for men being 37 years and for women being 32 years.

**Table 6.** Parameters for the fits to the number of men (M) and women (W) who are HIV positive among those suspected of having HIV (S) or whose HIV status is unknown (U).

|  | MS | MU | WS | WU |
| --- | --- | --- | --- | --- |
| Proportion positive | 0.59 | 0.074 | 0.51 | 0.104 |
| Norm | 869 | 1943 | 821 | 1896 |

Table 6 shows that 59% of the men and 51% of the women who are suspected of having HIV are HIV positive while 93% (100%-7%) of the men and 90% (100%-10%) of the women who are thought to be uninfected are HIV negative.

We can now calculate the proportion of false positives and false negatives if we assume that those that that are suspected of being positive are in fact positive and those whose status is unknown are in fact negative with the results given in Table 7. For example, if we assume that all those that are suspected of having HIV do in fact have HIV the number of false positive men will be (1-0.59)×869 = 356 which amounts to 356/(869+3672) = 7.8% of those assigned to the HIV-positive category.

**Table 7.** The proportion of men and women that are misclassified if ‘unknowns’ are treated as negative and ‘suspected’ are treated as positive.

|  | MP | MN | WP | WN |
| --- | --- | --- | --- | --- |
| Prop. misclassified | 0.078 | 0.051 | 0.082 | 0.072 |

Finally, we estimate the effect that this misclassification will have on the odds for being HIV-positive. For the odds in men the numerator will be increased by 7.8% and the denominator will be increased by 5.1% giving an overall increase in the odds ratio of 2.7% (1.078/1.051-1) while for women the increases will be 8.2% and 7.2% giving an overall increase in the odds ratio of 0.9%.

### Appendix 4: Attributable fractions

The attributable fractions for each AIDS-related condition were calculated as follows. Let the number of deaths in the control group (C) and each disease group (D) that are HIV-positive and negative be as indicated in Table 8.

**Table 8.** The number of deaths in the control group (C) and a given disease group (D) that are HIV positive and HIV-negative.

|  | HIV– | HIV+ |
| --- | --- | --- |
| C | $a$ | $b$ |
| D | $c$ | $d$ |

The odds-ratio for each condition D compared to the controls, C, is

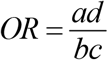

If HIV had no effect on the probability of dying of a particular condition the expected number of HIV-positive deaths from condition D would be

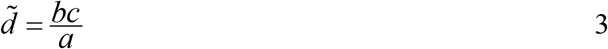

The HIV-attributable fraction of deaths in HIV-positive people with condition D is

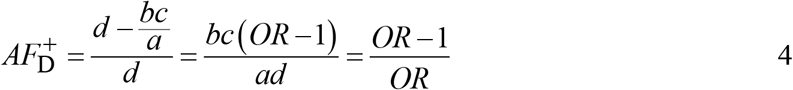

If 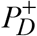 is the prevalence of HIV in those with condition D the disease attributable fraction of deaths in all those with condition D is

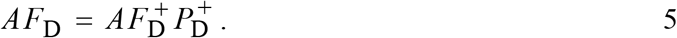

If *P*_D_ is the prevalence of condition D in the whole sample the population-attributable fraction for HIV due to condition *D* in the whole sample, is

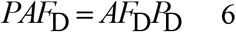

Finally, the attributable fraction for all AIDS-related conditions is the sum over Equation 6 for all AIDS-related conditions, D.

### Appendix 5: Daily mortality

By definition everyone in the sample had died and it is of interest to record the distribution of time between admission and deaths as shown in Figure 2. Mortality after admission was high with a median survival of 4 days and a mean survival of 6.7 days. On the day of admission 10% died, by the next day 27% had died and by the second day after admission 37% had died. By day sixteen 90% had died and by day forty 99% had died.

**Figure 2.**
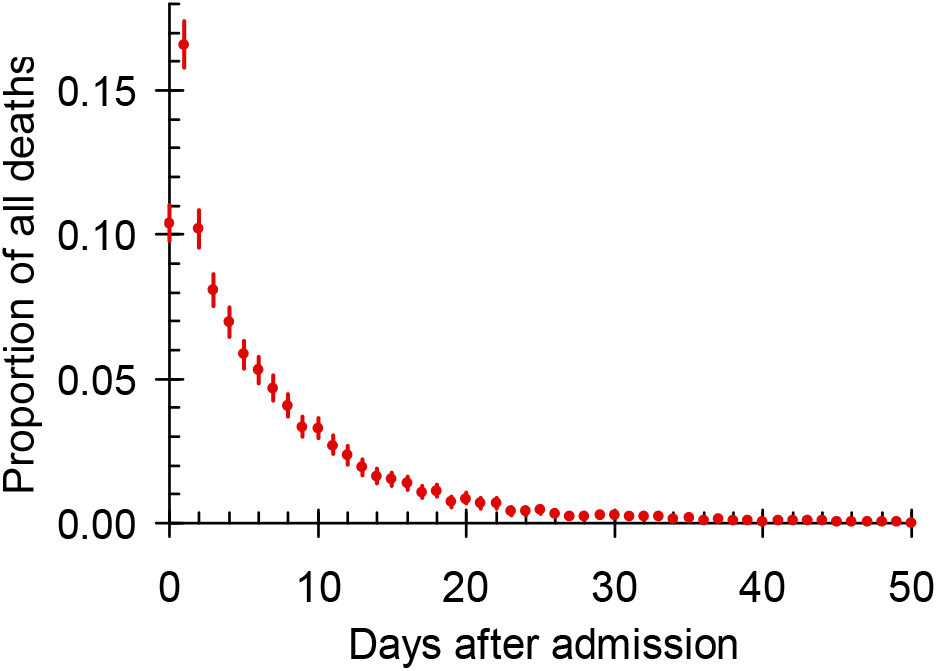
Deaths on a given number of days after admission on day zero as a proportion of all deaths.

### Appendix 6. Comparison with ANC data for Johannesburg Municipality

The prevalence of HIV in women attending antenatal clinics in the Johannesburg Municipality for the years 2006 to 2009^1–4^ is significantly less than in the control group used in this study with a peak prevalence at age 30 years of 42% as compared to 68% in the control group. Comparing the two prevalence estimates for those ages where ANC data are available gives an odds ratio for the control group in this study relative to the data from the ante-natal clinic surveys of 3.2 ± 0.7. Figure 3 shows a comparison of the two sets of data after rescaling the ANC data assuming an odds-ratio of 3.2.^1–4^

**Figure 3.**
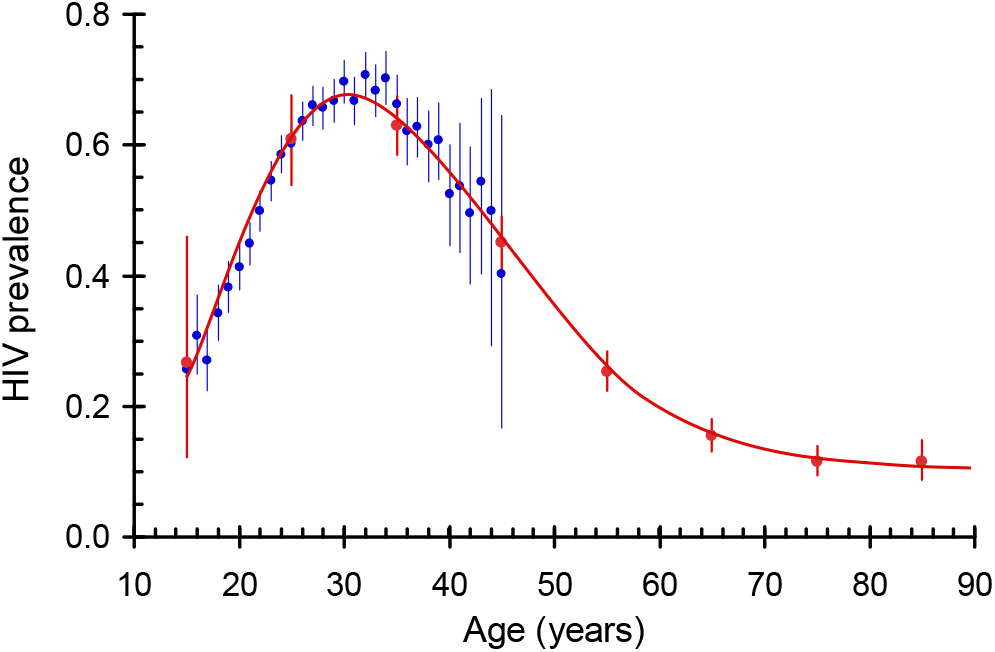
The prevalence of HIV as a function of age in the control group (red) and in the ANC data from Johannesburg Municipality (blue) for the years 2006 to 2009. The ANC data are scaled assuming an odds-ratio of 3.2.

We then recalculated the DAFs and the PAFS for each condition, after increasing with the results shown in Figure 4. For cryptococcosis, Kaposi’s sarcoma and *Pneumocystis carinii* the OR is very large so that *AF*^+^_*D*_ ≈ 1 and increasing the OR further will have little effect on the DAFs or PAFs. For respiratory conditions the OR is low, increasing the OR leads to a significant increase in the DAF and since the prevalence is high there will also be a significant increase in the PAF. For digestive conditions the OR is again low and increasing the OR leads to a significant increase in the DAF but the prevalence of digestive conditions is low so that the PAF remains low.

**Figure 4.**
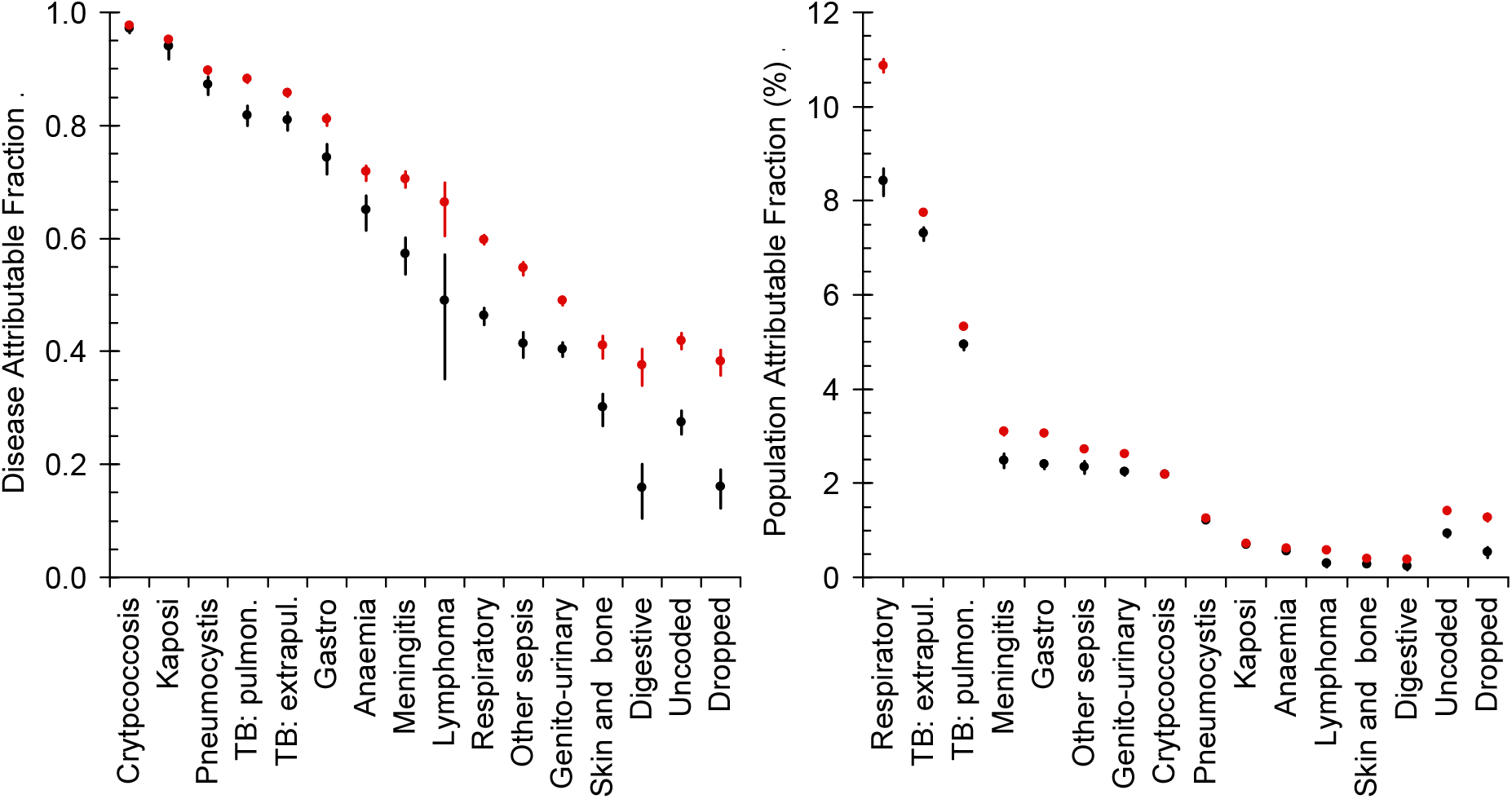
A: The disease-attributable fraction (DAF). B: The population-attributable fraction (PAF). Black: using the age-specific prevalence in the control group as described in the paper. Red: with the odds-ratios for each condition increased by a factor of 3.2.

### Appendix 7. Prevalence, odds and odds ratios for each disease class

Table 9 to Table 21 give by age, for each category in Table 3, the number of people that were HIV-positive (P+S), HIV-negative (N+U), the prevalence of HIV and the Odds for being HIV-positive with 95% confidence limits.

**Table 9.**
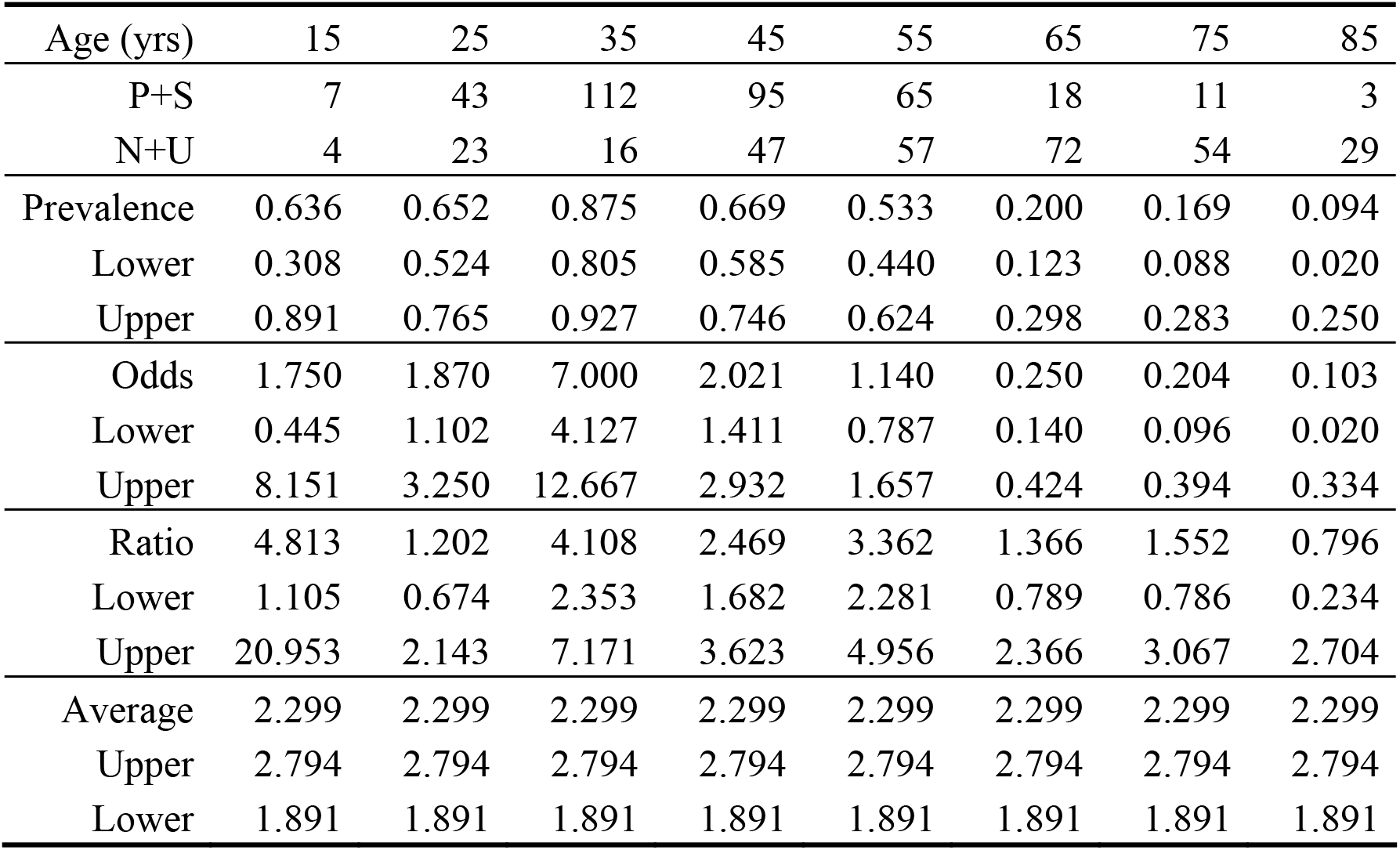
Not coded. Deaths that were not assigned an ICD-10 code.

**Table 10.** Dropped: A02-A07; A32-A39; A49-A87; B01-B18; B37-B43; B46; B50-B58; D01-D46; H52-H70; O90; P20

| Age (yrs) | 15 | 25 | 35 | 45 | 55 | 65 | 75 | 85 |
| --- | --- | --- | --- | --- | --- | --- | --- | --- |
| P+S | 4 | 47 | 79 | 51 | 33 | 10 | 7 | 3 |
| N+U | 6 | 25 | 27 | 41 | 63 | 32 | 39 | 31 |
| Prevalence | 0.400 | 0.653 | 0.745 | 0.554 | 0.344 | 0.238 | 0.152 | 0.088 |
| Lower | 0.122 | 0.531 | 0.651 | 0.447 | 0.250 | 0.121 | 0.063 | 0.019 |
| Upper | 0.738 | 0.761 | 0.825 | 0.658 | 0.448 | 0.395 | 0.289 | 0.237 |
| Odds | 0.667 | 1.880 | 2.926 | 1.244 | 0.524 | 0.313 | 0.179 | 0.097 |
| Lower | 0.138 | 1.134 | 1.869 | 0.808 | 0.333 | 0.137 | 0.068 | 0.019 |
| Upper | 2.811 | 3.187 | 4.713 | 1.925 | 0.810 | 0.652 | 0.406 | 0.310 |
| Ratio | 1.833 | 1.209 | 1.717 | 1.519 | 1.545 | 1.708 | 1.368 | 0.744 |
| Lower | 0.408 | 0.691 | 1.066 | 0.978 | 0.986 | 0.820 | 0.596 | 0.220 |
| Upper | 8.231 | 2.115 | 2.765 | 2.360 | 2.419 | 3.559 | 3.140 | 2.521 |
| Average | 1.489 | 1.489 | 1.489 | 1.489 | 1.489 | 1.489 | 1.489 | 1.489 |
| Upper | 1.842 | 1.842 | 1.842 | 1.842 | 1.842 | 1.842 | 1.842 | 1.842 |
| Lower | 1.204 | 1.204 | 1.204 | 1.204 | 1.204 | 1.204 | 1.204 | 1.204 |

**Table 11.** Controls: Malignant neo-plasms excluding Kaposi’s sarcoma, Hodgkin’s and Non-Hodgkin’s; disorders involving the immune mechanism; mental; other nervous system; heart and stroke; injury. C02-C45; C59-C80; C83; C90-C96; E03-E88; F01-F30; G10-G99; IU05-I95; Q24-Z91

| Age (yrs) | 15 | 25 | 35 | 45 | 55 | 65 | 75 | 85 |
| --- | --- | --- | --- | --- | --- | --- | --- | --- |
| P+S | 0 | 8 | 126 | 288 | 280 | 214 | 131 | 100 |
| N+U | 1 | 22 | 81 | 169 | 342 | 631 | 716 | 762 |
| Prevalence | 0.000 | 0.267 | 0.609 | 0.630 | 0.450 | 0.253 | 0.155 | 0.116 |
| Lower | 0.000 | 0.123 | 0.539 | 0.584 | 0.411 | 0.224 | 0.131 | 0.095 |
| Upper | 0.010 | 0.459 | 0.676 | 0.675 | 0.490 | 0.284 | 0.181 | 0.139 |
| Odds | 0.000 | 0.364 | 1.556 | 1.704 | 0.819 | 0.339 | 0.183 | 0.131 |
| Lower | 0.000 | 0.140 | 1.167 | 1.404 | 0.697 | 0.289 | 0.151 | 0.105 |
| Upper | 0.010 | 0.848 | 2.083 | 2.073 | 0.962 | 0.397 | 0.221 | 0.162 |

**Table 12.**
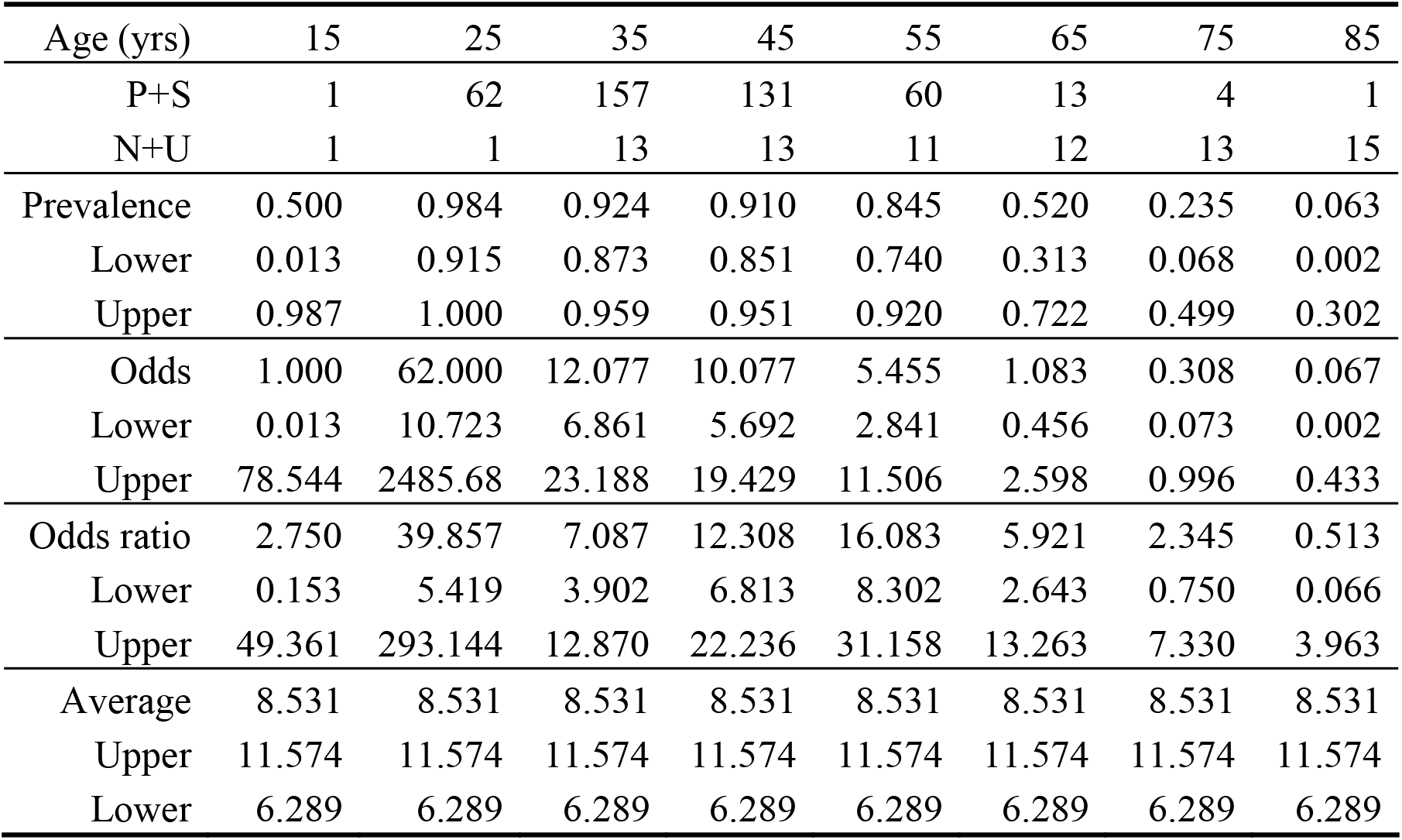
Infectious gastroenteritis and colitis, unspecified: A09

| Age (yrs) | 15 | 25 | 35 | 45 | 55 | 65 | 75 | 85 |
| --- | --- | --- | --- | --- | --- | --- | --- | --- |
| P+S | 1 | 62 | 157 | 131 | 60 | 13 | 4 | 1 |
| N+U | 1 | 1 | 13 | 13 | 11 | 12 | 13 | 15 |
| Prevalence | 0.500 | 0.984 | 0.924 | 0.910 | 0.845 | 0.520 | 0.235 | 0.063 |
| Lower | 0.013 | 0.915 | 0.873 | 0.851 | 0.740 | 0.313 | 0.068 | 0.002 |
| Upper | 0.987 | 1.000 | 0.959 | 0.951 | 0.920 | 0.722 | 0.499 | 0.302 |
| Odds | 1.000 | 62.000 | 12.077 | 10.077 | 5.455 | 1.083 | 0.308 | 0.067 |
| Lower | 0.013 | 10.723 | 6.861 | 5.692 | 2.841 | 0.456 | 0.073 | 0.002 |
| Upper | 78.544 | 2485.68 | 23.188 | 19.429 | 11.506 | 2.598 | 0.996 | 0.433 |
| Odds ratio | 2.750 | 39.857 | 7.087 | 12.308 | 16.083 | 5.921 | 2.345 | 0.513 |
| Lower | 0.153 | 5.419 | 3.902 | 6.813 | 8.302 | 2.643 | 0.750 | 0.066 |
| Upper | 49.361 | 293.144 | 12.870 | 22.236 | 31.158 | 13.263 | 7.330 | 3.963 |
| Average | 8.531 | 8.531 | 8.531 | 8.531 | 8.531 | 8.531 | 8.531 | 8.531 |
| Upper | 11.574 | 11.574 | 11.574 | 11.574 | 11.574 | 11.574 | 11.574 | 11.574 |
| Lower | 6.289 | 6.289 | 6.289 | 6.289 | 6.289 | 6.289 | 6.289 | 6.289 |

**Table 13.** TB: pulmonary A15; A16

| Age (yrs) | 15 | 25 | 35 | 45 | 55 | 65 | 75 | 85 |
| --- | --- | --- | --- | --- | --- | --- | --- | --- |
| P+S | 14 | 206 | 494 | 330 | 174 | 30 | 5 | 0 |
| N+U | 4 | 19 | 24 | 33 | 45 | 32 | 9 | 2 |
| Prevalence | 0.778 | 0.916 | 0.954 | 0.909 | 0.795 | 0.484 | 0.357 |  |
| Lower | 0.524 | 0.871 | 0.932 | 0.875 | 0.735 | 0.355 | 0.128 |  |
| Upper | 0.936 | 0.948 | 0.970 | 0.937 | 0.846 | 0.614 | 0.649 |  |
| Odds | 3.500 | 10.842 | 20.583 | 10.000 | 3.867 | 0.938 | 0.556 |  |
| Lower | 1.099 | 6.768 | 13.672 | 6.981 | 2.772 | 0.550 | 0.146 |  |
| Upper | 14.602 | 18.377 | 32.435 | 14.771 | 5.493 | 1.593 | 1.846 |  |
| Odds ratio | 0.000 | 6.970 | 12.078 | 12.214 | 11.401 | 5.124 | 4.233 |  |
| Lower | 0.000 | 4.035 | 7.689 | 8.260 | 7.934 | 3.011 | 1.391 |  |
| Upper | 0.000 | 12.039 | 18.972 | 18.061 | 16.384 | 8.721 | 12.883 |  |
| Average | 9.620 | 9.620 | 9.620 | 9.620 | 9.620 | 9.620 | 9.620 |  |
| Upper | 11.672 | 11.672 | 11.672 | 11.672 | 11.672 | 11.672 | 11.672 |  |
| Lower | 7.928 | 7.928 | 7.928 | 7.928 | 7.928 | 7.928 | 7.928 |  |

**Table 14.** TB: extra-pulmonary: A17-A19

| Age (yrs) | 15 | 25 | 35 | 45 | 55 | 65 | 75 | 85 |
| --- | --- | --- | --- | --- | --- | --- | --- | --- |
| P+S | 4 | 141 | 368 | 244 | 88 | 17 | 3 | 2 |
| N+U | 1 | 10 | 20 | 23 | 17 | 6 | 3 | 2 |
| Prevalence | 0.800 | 0.934 | 0.948 | 0.914 | 0.838 | 0.739 | 0.500 | 0.500 |
| Lower | 0.284 | 0.882 | 0.922 | 0.874 | 0.753 | 0.516 | 0.118 | 0.068 |
| Upper | 0.995 | 0.968 | 0.968 | 0.945 | 0.903 | 0.898 | 0.882 | 0.932 |
| Odds | 4.000 | 14.100 | 18.400 | 10.609 | 5.176 | 2.833 | 1.000 | 1.000 |
| Lower | 0.396 | 7.445 | 11.741 | 6.908 | 3.056 | 1.066 | 0.134 | 0.072 |
| Upper | 196.973 | 30.044 | 30.480 | 17.053 | 9.284 | 8.776 | 7.466 | 13.796 |
| Odds ratio | 11.000 | 9.064 | 10.797 | 12.958 | 15.263 | 15.486 | 7.620 | 7.692 |
| Lower | 1.064 | 4.503 | 6.625 | 8.215 | 8.878 | 5.994 | 1.517 | 1.061 |
| Upper | 113.736 | 18.244 | 17.597 | 20.439 | 26.242 | 40.009 | 38.269 | 55.779 |
| Average | 12.112 | 12.112 | 12.112 | 12.112 | 12.112 | 12.112 | 12.112 | 12.112 |
| Upper | 15.506 | 15.506 | 15.506 | 15.506 | 15.506 | 15.506 | 15.506 | 15.506 |
| Lower | 9.461 | 9.461 | 9.461 | 9.461 | 9.461 | 9.461 | 9.461 | 9.461 |

**Table 15.** Other sepsis, Infectious and parasitic: A41

| Age (yrs) | 15 | 25 | 35 | 45 | 55 | 65 | 75 | 85 |
| --- | --- | --- | --- | --- | --- | --- | --- | --- |
| P+S | 6 | 84 | 184 | 154 | 73 | 26 | 9 | 4 |
| N+U | 3 | 17 | 25 | 49 | 47 | 81 | 69 | 46 |
| Prevalence | 0.667 | 0.832 | 0.880 | 0.759 | 0.608 | 0.243 | 0.115 | 0.080 |
| Lower | 0.299 | 0.744 | 0.829 | 0.694 | 0.515 | 0.165 | 0.054 | 0.022 |
| Upper | 0.925 | 0.899 | 0.921 | 0.816 | 0.696 | 0.335 | 0.208 | 0.192 |
| Odds | 2.000 | 4.941 | 7.360 | 3.143 | 1.553 | 0.321 | 0.130 | 0.087 |
| Lower | 0.427 | 2.910 | 4.831 | 2.265 | 1.062 | 0.198 | 0.057 | 0.023 |
| Upper | 12.359 | 8.880 | 11.670 | 4.428 | 2.291 | 0.505 | 0.262 | 0.238 |
| Odds ratio | 5.500 | 3.176 | 4.319 | 3.839 | 4.580 | 1.754 | 0.994 | 0.669 |
| Lower | 1.105 | 1.759 | 2.729 | 2.683 | 3.076 | 1.086 | 0.481 | 0.231 |
| Upper | 27.375 | 5.738 | 6.834 | 5.492 | 6.818 | 2.834 | 2.053 | 1.934 |
| Average | 3.100 | 3.100 | 3.100 | 3.100 | 3.100 | 3.100 | 3.100 | 3.100 |
| Upper | 3.729 | 3.729 | 3.729 | 3.729 | 3.729 | 3.729 | 3.729 | 3.729 |
| Lower | 2.576 | 2.576 | 2.576 | 2.576 | 2.576 | 2.576 | 2.576 | 2.576 |

**Table 16.**
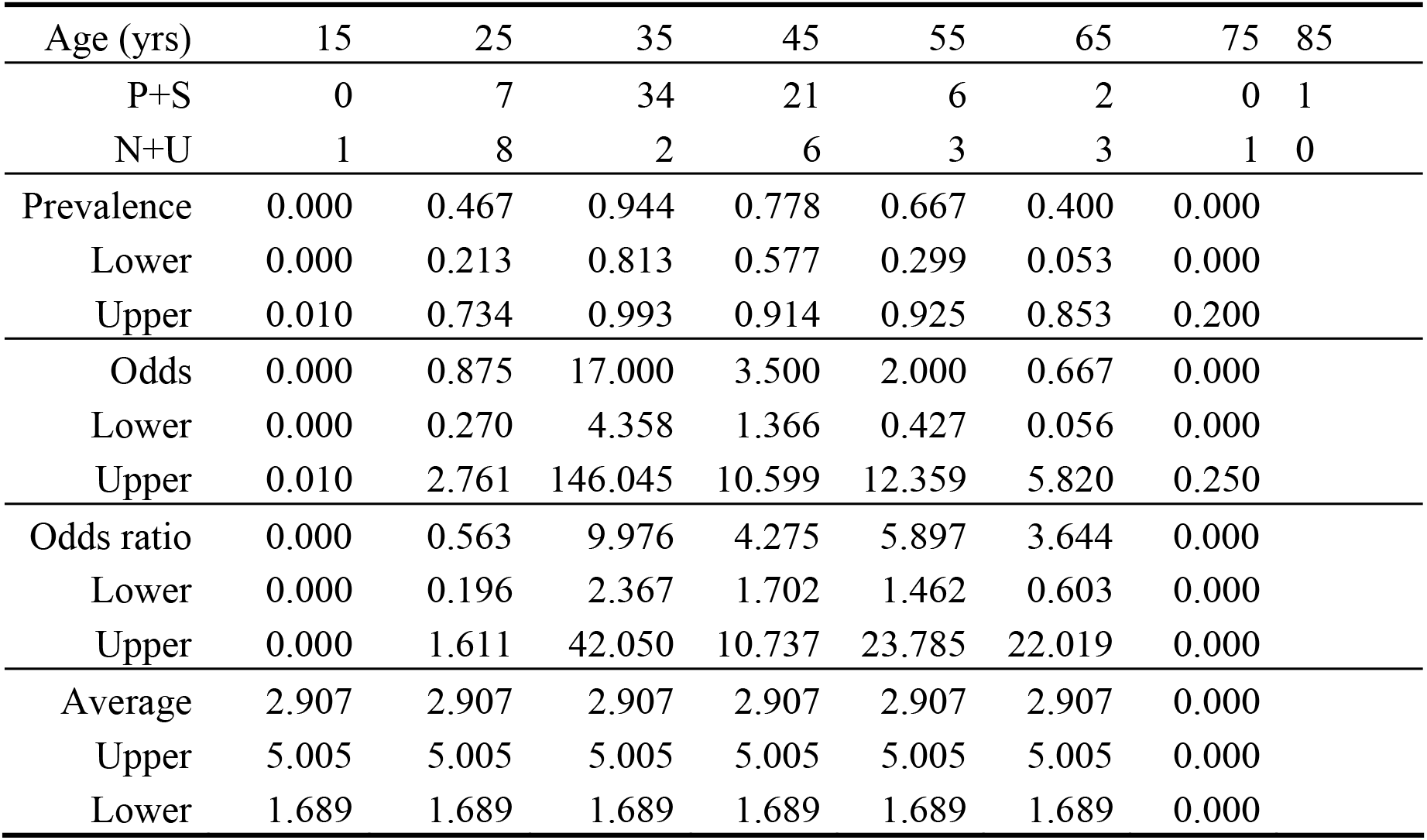
Hodgkin’s and non-Hodgkin’s lymphoma: C81; C85

**Table 17.**
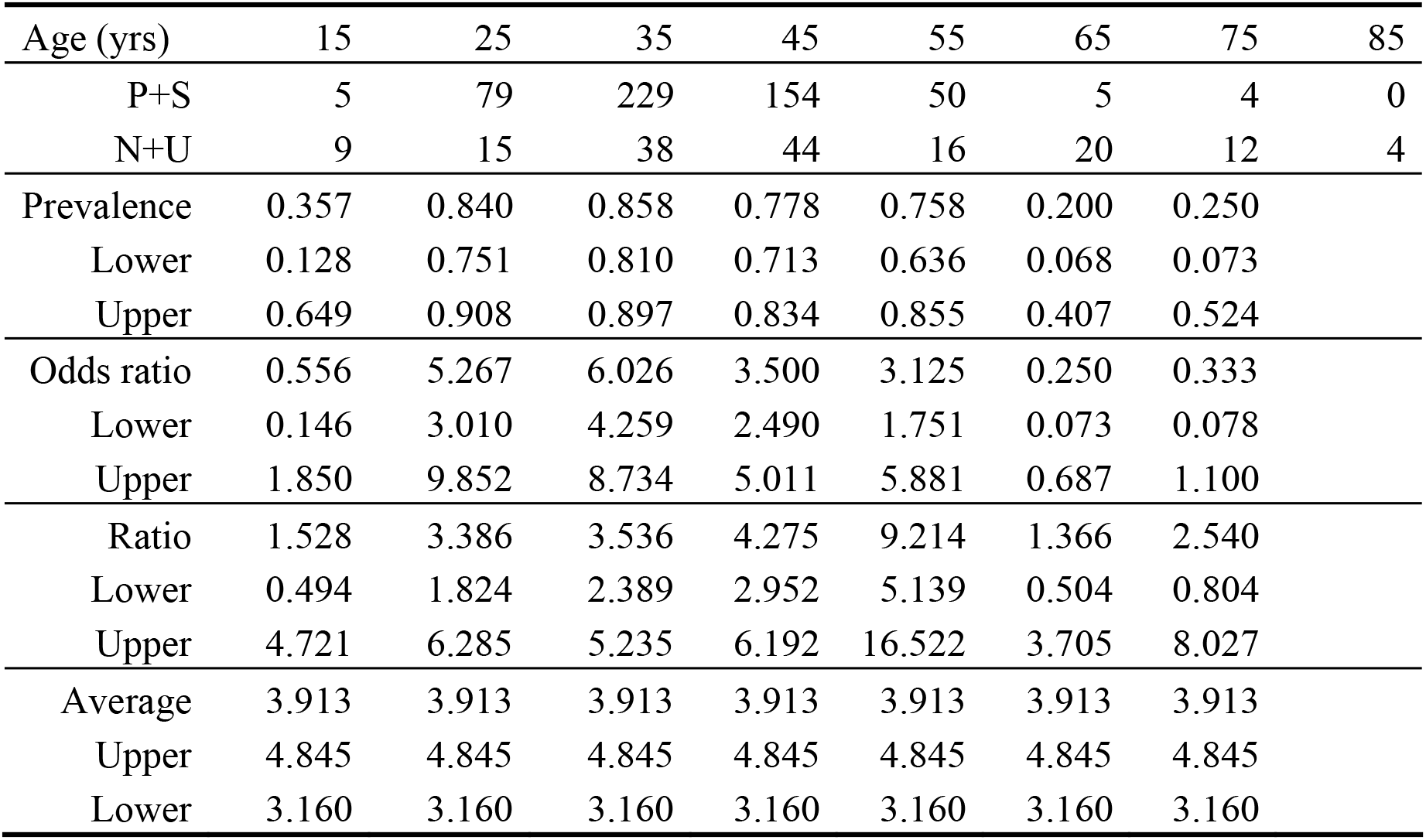
Meningitis: G00-G03

**Table 18.** Respiratory, Pneumonia, COPD; lower respiratory: J01-J99

| Age (yrs) | 15 | 25 | 35 | 45 | 55 | 65 | 75 | 85 |
| --- | --- | --- | --- | --- | --- | --- | --- | --- |
| P+S | 18 | 286 | 648 | 545 | 266 | 73 | 42 | 13 |
| N+U | 10 | 65 | 91 | 123 | 217 | 211 | 142 | 90 |
| Prevalence | 0.643 | 0.815 | 0.877 | 0.816 | 0.551 | 0.257 | 0.228 | 0.126 |
| Lower | 0.441 | 0.770 | 0.851 | 0.784 | 0.505 | 0.207 | 0.170 | 0.069 |
| Upper | 0.814 | 0.854 | 0.900 | 0.845 | 0.596 | 0.312 | 0.296 | 0.206 |
| Odds ratio | 1.800 | 4.400 | 7.121 | 4.431 | 1.226 | 0.346 | 0.296 | 0.144 |
| Lower | 0.788 | 3.351 | 5.710 | 3.637 | 1.021 | 0.261 | 0.204 | 0.074 |
| Upper | 4.365 | 5.853 | 8.969 | 5.433 | 1.473 | 0.453 | 0.420 | 0.260 |
| Ratio | 4.950 | 2.829 | 4.179 | 5.412 | 3.614 | 1.891 | 2.254 | 1.111 |
| Lower | 1.617 | 1.919 | 3.126 | 4.209 | 2.852 | 1.366 | 1.524 | 0.603 |
| Upper | 15.157 | 4.169 | 5.585 | 6.959 | 4.581 | 2.617 | 3.334 | 2.046 |
| Average | 3.322 | 3.322 | 3.322 | 3.322 | 3.322 | 3.322 | 3.322 | 3.322 |
| Upper | 3.738 | 3.738 | 3.738 | 3.738 | 3.738 | 3.738 | 3.738 | 3.738 |
| Lower | 2.952 | 2.952 | 2.952 | 2.952 | 2.952 | 2.952 | 2.952 | 2.952 |

**Table 19.**
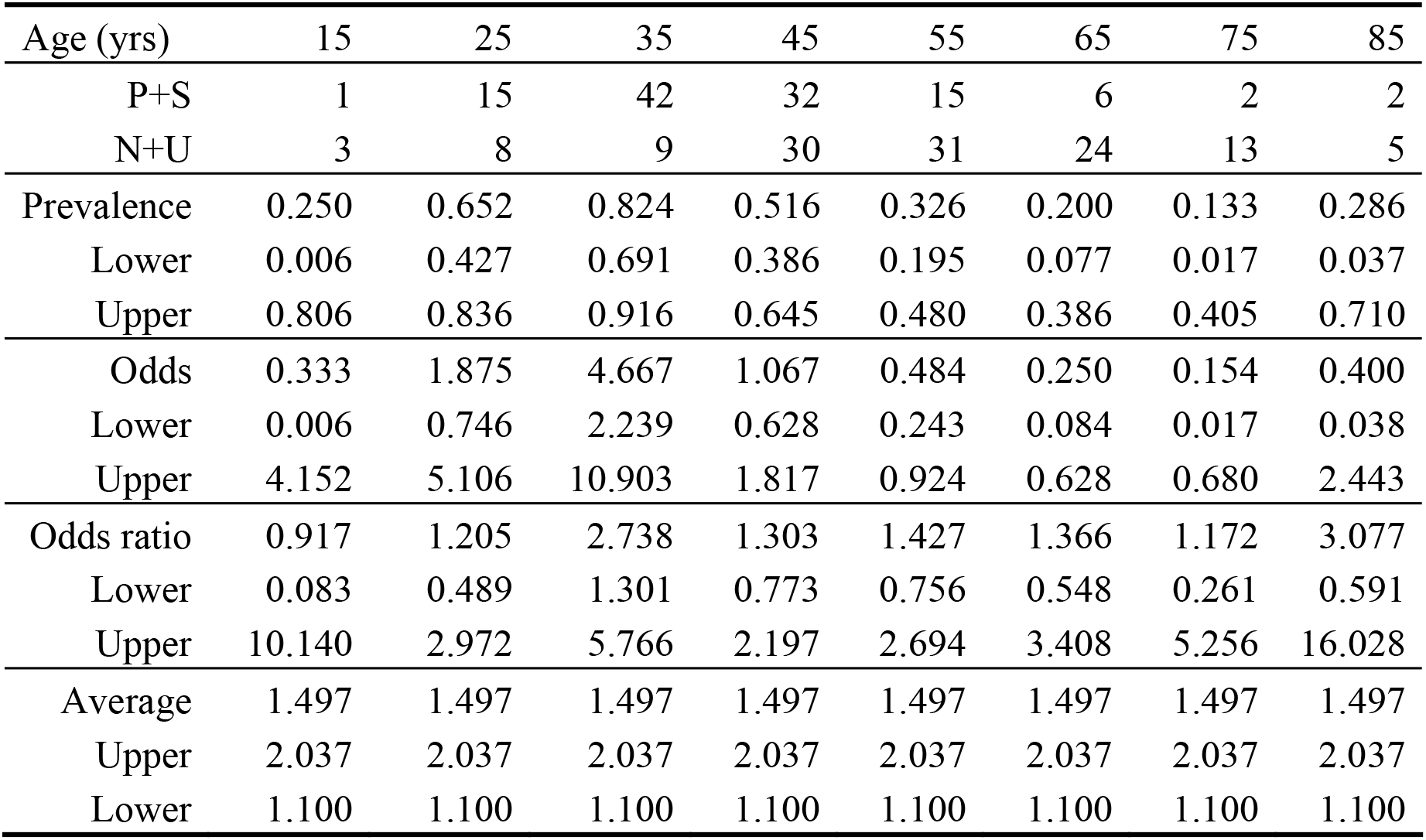
Digestive, Liver: K25-K95

**Table 20.**
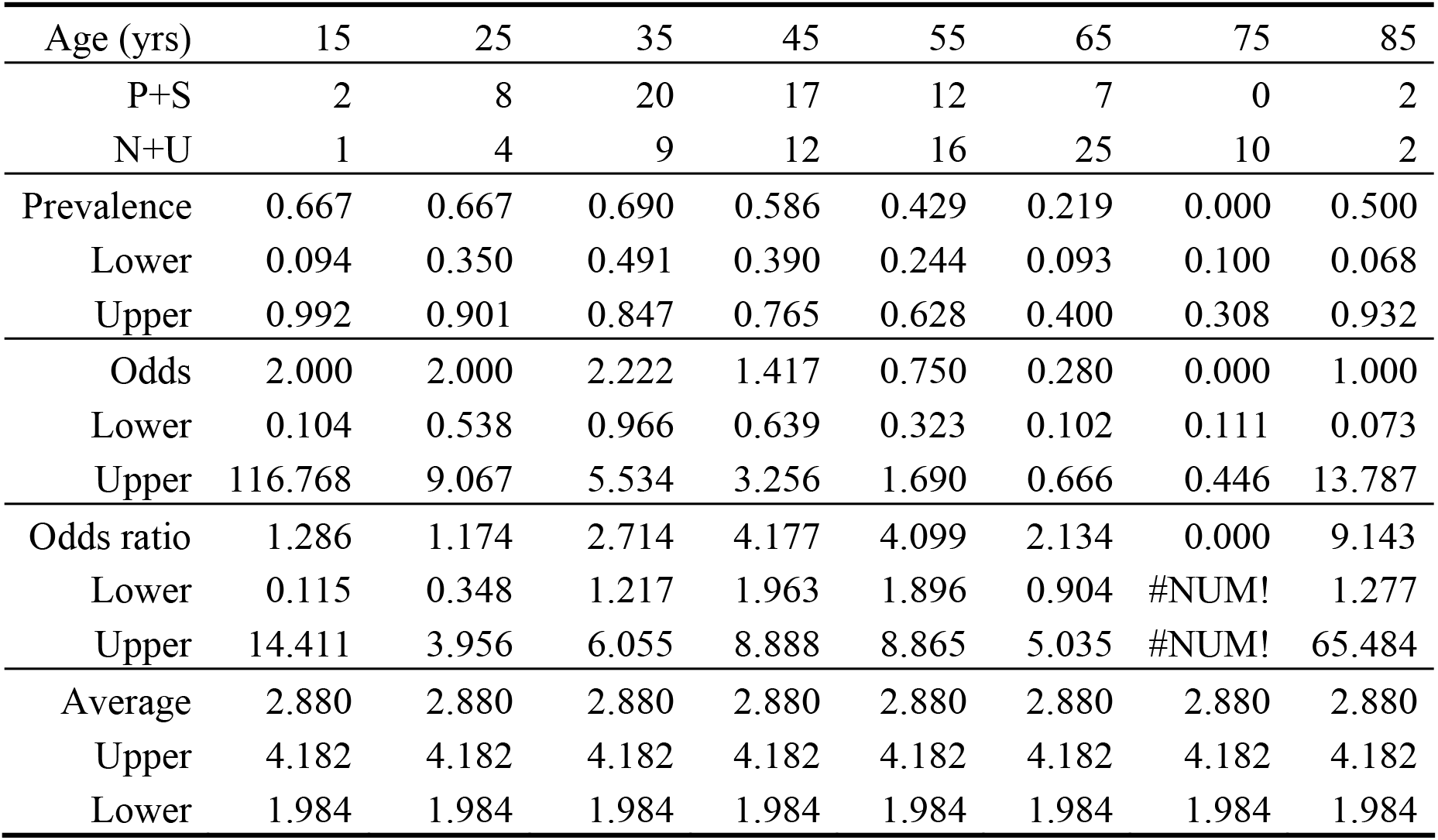
Skin and bone: L03-L95; M01-M99

**Table 21.** Genito-urinary infections: N00-N92.

| Age (yrs) | 15 | 25 | 35 | 45 | 55 | 65 | 75 | 85 |
| --- | --- | --- | --- | --- | --- | --- | --- | --- |
| P+S | 2 | 57 | 130 | 153 | 70 | 26 | 17 | 9 |
| N+U | 7 | 20 | 28 | 51 | 90 | 100 | 61 | 45 |
| Prevalence | 0.222 | 0.740 | 0.823 | 0.750 | 0.438 | 0.206 | 0.218 | 0.167 |
| Lower | 0.028 | 0.628 | 0.754 | 0.685 | 0.359 | 0.139 | 0.132 | 0.079 |
| Upper | 0.600 | 0.834 | 0.879 | 0.808 | 0.518 | 0.288 | 0.326 | 0.293 |
| Odds ratio | 0.286 | 2.850 | 4.643 | 3.000 | 0.778 | 0.260 | 0.279 | 0.200 |
| Lower | 0.029 | 1.686 | 3.068 | 2.172 | 0.561 | 0.162 | 0.153 | 0.086 |
| Upper | 1.501 | 5.008 | 7.257 | 4.204 | 1.075 | 0.404 | 0.483 | 0.414 |
| Ratio | 0.184 | 1.672 | 5.671 | 8.846 | 4.251 | 1.981 | 2.144 | 1.829 |
| Lower | 0.037 | 0.971 | 3.660 | 6.216 | 2.955 | 1.238 | 1.214 | 0.873 |
| Upper | 0.906 | 2.880 | 8.786 | 12.588 | 6.116 | 3.170 | 3.787 | 3.831 |
| Average | 4.185 | 4.185 | 4.185 | 4.185 | 4.185 | 4.185 | 4.185 | 4.185 |
| Upper | 5.037 | 5.037 | 5.037 | 5.037 | 5.037 | 5.037 | 5.037 | 5.037 |
| Lower | 3.477 | 3.477 | 3.477 | 3.477 | 3.477 | 3.477 | 3.477 | 3.477 |

The data in Table 9 to Table 21 are plotted against age with the overall estimate of the OR and 95% confidence limits. The confidence limits on the average values are calculated using only the binomial sampling errors. For genitor-urinary conditions there is further variability over and above the sampling errors.

**Figure 5.**
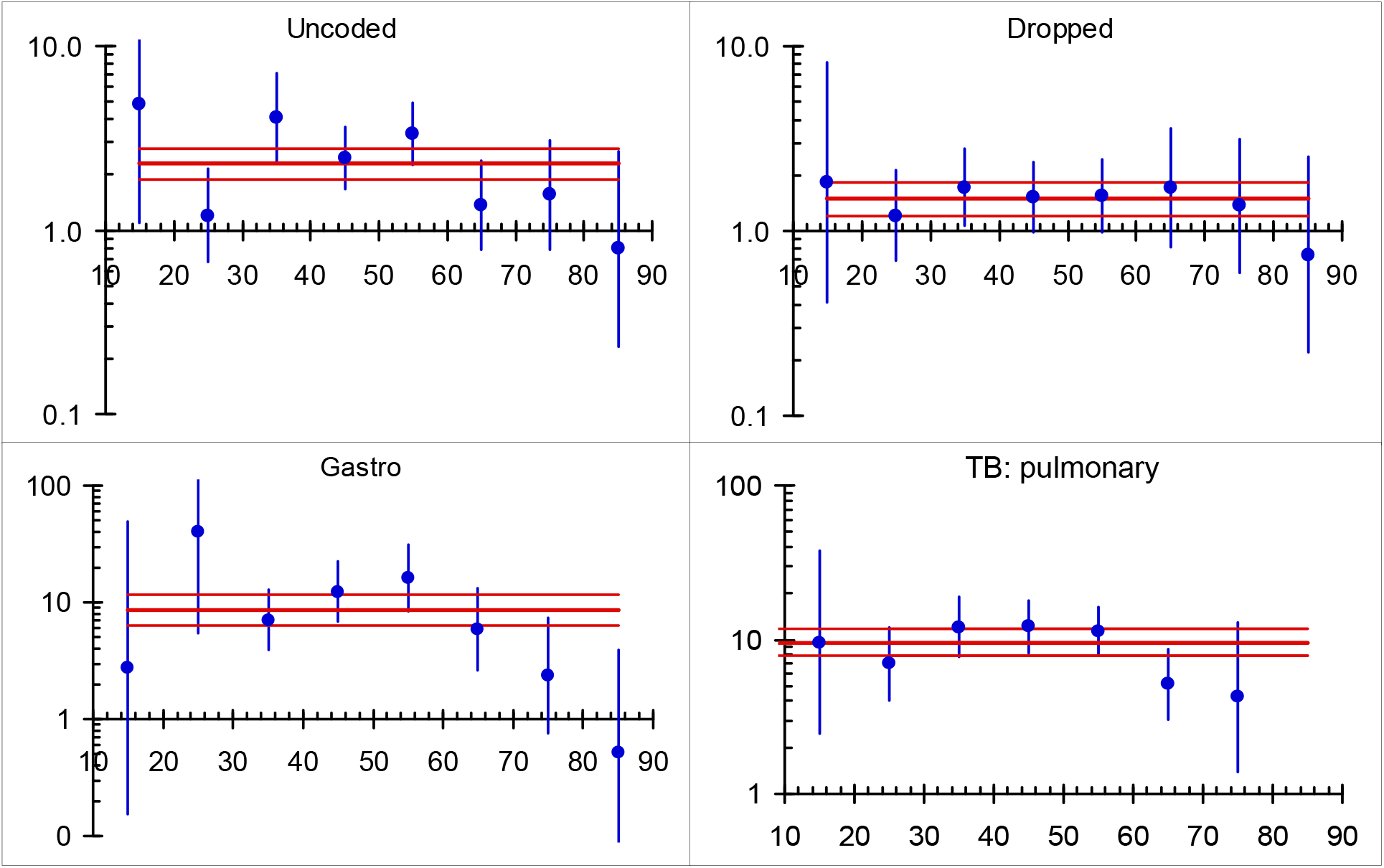
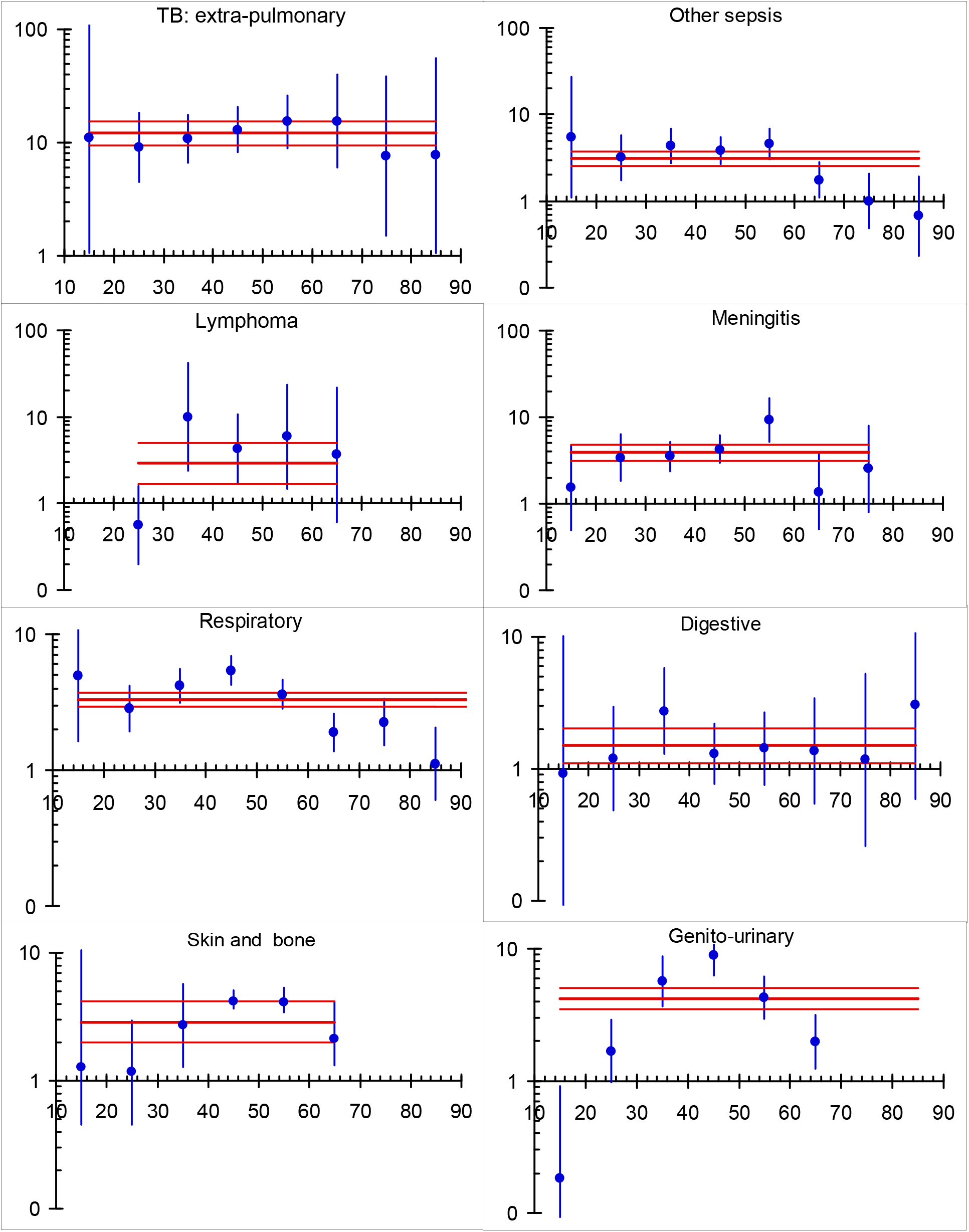
Odds-ratios by age for each of the conditions noted on the figure with the average values and 95% confidence limits for the points and for the average value.

### Appendix 8. Age-standardization for small numbers

Age-standardization is important as the prevalence of HIV is very high among those less than 50 years of age but much lower among those more than 50 years of age as seen in Figure 5. For certain conditions in which the prevalence of HIV was high, so that the number that were HIV-negative was very small it was not possible to standardize on age. We therefore compared the odds ratios with and without standardization for those conditions for which age-standardization could be done reliably using the data in Table 11 to Table 21 with the result shown in Figure 6. The relationship between the age-standardized odds ratio, *A*, and the crude odds ratio, *C*, is

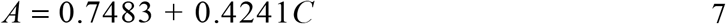

and we used this relationship to estimate the adjusted odds ratios for pulmonary cryptococcus, Kaposi’s sarcoma, *Pneumocystis carinii,* and diseases of the blood and blood forming organs.

**Figure 6.**
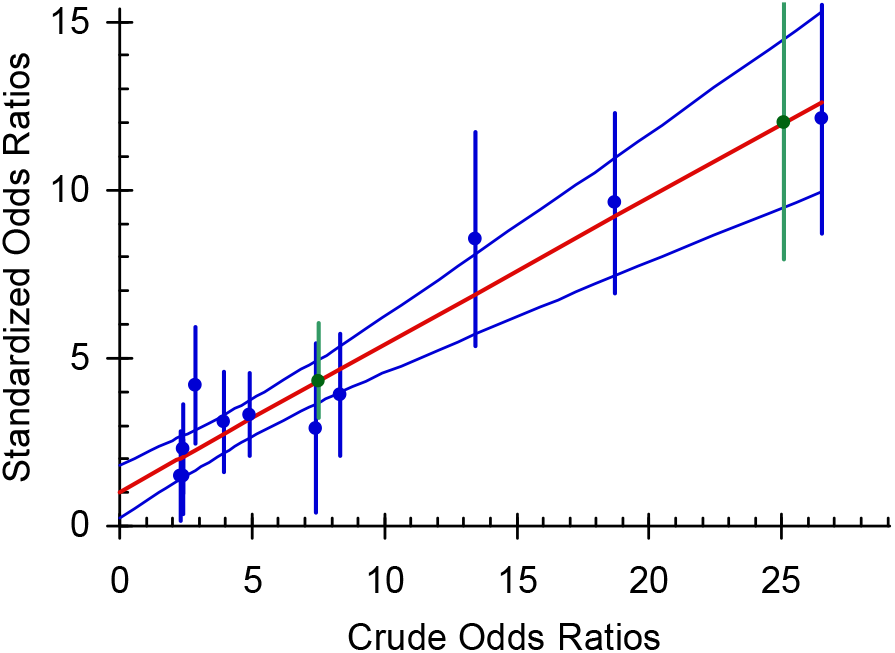
Odds ratios standardized for age plotted against crude odds ratios. Blue points: standardized odds ratio measured; Green points: standardized odds ratios for diseases of the blood and blood forming organs and for *Pneumocystis carinii* calculated from the fitted line.

